# Network communication models narrow the gap between the modular organization of structural and functional brain networks

**DOI:** 10.1101/2022.02.18.480871

**Authors:** Caio Seguin, Sina Mansour L, Olaf Sporns, Andrew Zalesky, Fernando Calamante

## Abstract

Structural and functional brain networks are modular. Canonical functional systems, such as the default mode network, are well-known modules of the human brain and have been implicated in a large number of cognitive, behavioral and clinical processes. However, modules delineated in structural brain networks inferred from tractography generally do not recapitulate canonical functional systems. Neuroimaging evidence suggests that functional connectivity between regions in the same systems is not always underpinned by anatomical connections. As such, direct structural connectivity alone would be insufficient to characterize the functional modular organization of the brain. Here, we demonstrate that augmenting structural brain networks with models of indirect (polysynaptic) communication unveils a modular network architecture that more closely resembles the brain’s established functional systems. We find that diffusion models of polysynaptic connectivity, particularly communicability, narrow the gap between the modular organization of structural and functional brain networks by 20–60%, whereas routing models based on single efficient paths do not improve mesoscopic structure-function correspondence. This suggests that functional modules emerge from the constraints imposed by local network structure that facilitates diffusive neural communication. Our work establishes the importance of modeling polysynaptic communication to understand the structural basis of functional systems.

## INTRODUCTION

The human brain is a complex network of interconnected neural elements [1, 2]. Using magnetic resonance imaging (MRI), connectivity between brain areas can be mapped in terms of structural links denoting anatomical white matter connections, or functional links capturing statistical patterns of co-activation over time [3]. Understanding the interplay between these two modalities of brain connectivity, i.e., how physical connections constrain and facilitate synchronized interregional activity, is a central challenge in modern neuroscience [4–6].

The presence of a modular architecture is a hallmark of both structural and functional human brain networks [7, 8]. Regions within the same module tend to be densely and strongly interconnected, while connectivity between regions in different modules is typically sparse and weak. Modular structure provides a mesoscopic account of network organization that is poised between local and global topological properties [9]. A large body of knowledge, comprising studies of both structural connectivity (SC) and functional connectivity (FC), indicates the brain’s modular architecture plays an important role in development [10], aging [11, 12], learning [13] and cognitive performance [14, 15], as well as in a range of mental health and neurodegenerative conditions [16].

Interestingly, however, there is a weak correspondence between the modular organization of structural and functional networks [6, 9, 17]. Structural modules are spatially compact and contiguous. With the exception of reports of homotopic modules located along the medial wall [18], structural modules are usually restricted to a single hemisphere. This modular structure is conjectured to reduce wiring costs associated with the creation and maintenance of long-range physical connections [19], while increasing network resilience by restricting the flow of pathological agents and maladaptive perturbations from their loci of origin [16]. In contrast, functional networks are characterized by spatially distributed modules, which often comprise distant and homotopic regions [20, 21]. Functional modules obtained through community detection methods closely recapitulate canonical brain systems and intrinsic resting-state networks identified through alternative data-driven techniques [22, 23] and meta-analyses of task-evoked activity [24]. This functional modular architecture is thought to promote segregated clusters of specialized information processing that correspond to specific cognitive domains [7, 8, 22].

The mismatch between structural and functional modules is most evident for multimodal brain systems involved in high-order cognition [6]. For instance, community detection methods applied to structural networks fail to identify the default mode or frontoparietal control networks, which can be retrieved following the application of the same methods to functional networks [20]. An explanation for this discordance comes from evidence that the FC between regions within the same functional systems is not entirely underpinned by direct anatomical connections [25]. For example, neuroimaging studies report weak to absent white matter connectivity between portions of the parietal cortex and the precuneus involved in the default mode network [26, 27]. As such, communication between certain functionally coupled—yet anatomically unconnected—regions must rely on polysynaptic signaling mediated by intermediate areas. SC alone would therefore provide an incomplete account of the mesoscale functional organization of the human brain [6].

Here, we hypothesize that network communication models can narrow the gap between structural and functional brain modules. This class of graph-theoretical models describes polysynaptic interactions between anatomically unconnected brain areas by modeling neural communication on top of SC [28, 29]. As such, these models can be used to augment SC—which encodes only direct anatomical connections—into communication matrices (CMs) that estimate interactions between all regions pairs in the brain [30]. Recent reports indicate that modeling polysynaptic communication improves predictions of FC [31], effective connectivity directionality [32], and individual variation in human behavior and cognition [30]. Based on this, we hypothesize that community detection applied to CMs will yield a modular architecture that more closely recapitulates the organization of the brain into canonical functional systems, compared to the modular structure of SC.

Importantly, it remains unclear which conceptualizations of polysynaptic transmission best describe largescale neural signaling. While numerous putative network communication models have been proposed [28], efforts to systematically compare and biologically validate different approaches have been limited. Here, we consider four popular measures of brain network communication, selected to cover a wide range of previously explored signaling strategies: i) shortest path efficiency [33, 34], ii) navigation efficiency [35, 36], iii) search information [31, 37], and iv) communicability [38, 39] (Table I). We conjecture that determining which models best account for the emergence of the brain’s mesoscopic functional organization will provide insight into the underlying mechanisms of large-scale neural signaling.

**TABLE I.** Summary of the four conceptualizations of neural communication investigated. Technical details are provided in the *Materials and Methods*.

| Network communication measure | Description |
| --- | --- |
| Shortest path efficiency (SPE) | Communication efficiency under the shortest path routing model. Communication occurs via optimally efficient paths. |
| Navigation efficiency (NE) | Communication efficiency under the navigation model. Communication occurs via geometrically greedy paths. |
| Search information (SI) | Amount of information required for a random walker to travel via shortest paths. Quantifies the accessibility of efficient paths in the network. |
| Communicability (CMY) | Weighted sum of all walk lengths between region pairs. Proposes communication via diffusive broadcasting. |

## RESULTS

Structural and functional connectivity were mapped for a sample for 1000 healthy adults participating in the Human Connectome Project (HCP; Materials and Methods) [40]. Connectivity data from individual participants were combined to generate the group-level structural and functional networks comprising 200 cortical regions defined in the Schaefer parcellation [41]. Following previous network modeling work [42, 43], we confined our analyses to intra-hemispheric networks (*N* = 100 regions for left and right hemispheres) to avoid well-known limitations in the mapping of cross-hemispheric fibers using tractography [44]. Network communication models were used to transform SC into communication matrices (CMs) capturing both direct and indirect interregional interactions under 4 putative models of polysynaptic signaling, thus yielding 4 different CMs (Fig 1a; Materials and Methods).

**FIG. 1.**
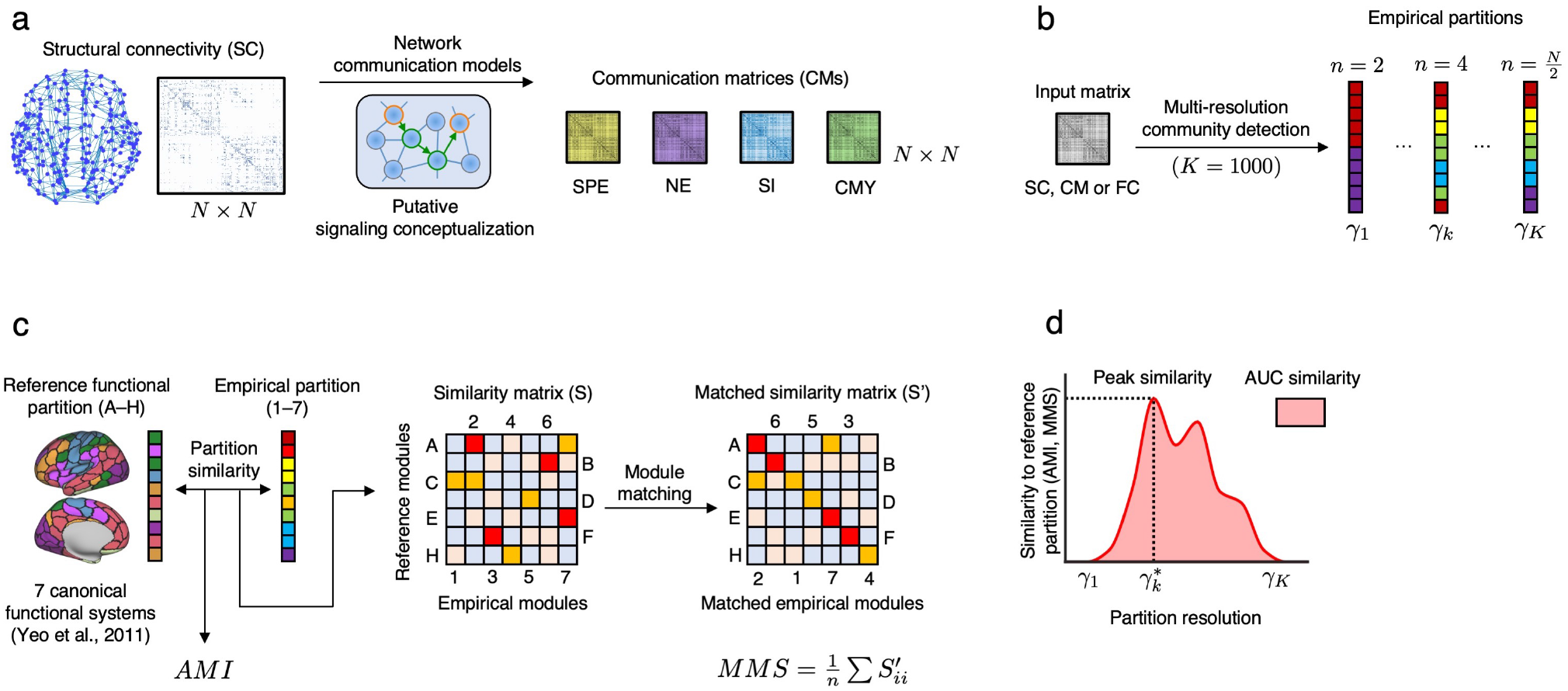
Methodology overview. **(a)** Network communication measures were used to model polysynaptic signaling on a group-level structural brain network. Four putative conceptualisations of neural signaling were considered: shortest path efficiency (SPE), navigation efficiency (NE), search information (SI) and communicability (CMY), resulting in a set of four communication matrices (CMs). While structural connectivity is sparse and only encodes direct anatomical connections, CMs are complete graphs and model both direct and polysynaptic interactions between regions. **(b)** Schematic of the multi-resolution community detection routine. For a given input matrix, a range of 1000 resolution parameters *γ* was selected to obtain partitions ranging from 2 to *N/*2 modules. **(c)** Schematics of partition similarity measures. The functional systems delineated by Yeo and colleagues [22] were adopted as the reference partition. Empirical and reference partitions were compared using the adjusted mutual information (AMI) and the mean matched similarity (MMS) measures. MMS is computed based on a one-to-one matching between reference and empirical modules. **(d)** The correspondence (MMS, AMI) between partitions was summarized using the peak and area-under-the-curve (AUC) similarity computed across multiple partition resolutions.

A well-established division of cortical regions into 7 canonical functional systems (also known as intrinsic resting-state networks) [22] was adopted as a reference partition of mesoscale functional organization. This partition has been consolidated as a reference frame for the study of hierarchical cortical organization [45, 46], large-scale neural dynamics [47], and brain-behavior relationships [48, 49], with mounting evidence supporting the cognitive and clinical relevance of these functional systems [50, 51].

SC, FC, and CMs were partitioned into modules using a multi-resolution community detection routine [52–54] based on the Louvain algorithm for modularity maximization [55] (Fig 1b; *Materials and Methods*). An important methodological challenge in the study of modularity is the need to select a resolution of modular decomposition. For any input network, the number of modules identified by the Louvain algorithm is expected to grow as a function of the resolution parameter *γ*. While a number of heuristics have been proposed to select *γ*, there is no consensus on best practices and the choice of resolution remains arbitrary in most applications [56]. To circumvent this issue, we considered modules obtained for 1000 values sampled across a meaningful portion of the *γ* parameter space. More specifically, we determined, separately for each input matrix, the range of *γ* values resulting in partitions ranging from 2 to *N/*2 modules. This yielded a multi-resolution set of partitions that allowed us to investigate the relation between structural and functional modules in a *γ*-resolved manner.

Empirical partitions obtained for CMs, SC and FC were then compared to the reference functional partition by means of two complementary measures: adjusted mutual information (AMI) and mean matched similarity (MMS) (Fig 1c; *Materials and Methods*). The AMI quantifies the amount of information shared between two partitions, while correcting for agreements solely due to chance [57]. The AMI provides a “global” assessment of partition similarly that does not capture the alignment between specific pairs of reference and empirical modules. To investigate which specific functional systems are best explained by structural modules, we developed a new measure called mean matched similarity (MMS). To compute MMS, we first calculate a similarity matrix that quantifies the agreement between all pairs of reference and empirical modules. The similarity matrix is used to perform an explicit one-to-one matching of reference to empirical modules by solving the assignment problem. This process can be thought of as a reshuffling of the columns (or rows) of the similarity matrix in order to maximize the sum of values along its main diagonal, i.e., maximize the total similarity of reference modules matched to empirical modules. Each entry along the main diagonal of the matched similarity matrix quantifies how well an individual reference functional system is captured by an empirical partition. MMS is defined as the mean of the matched matrix diagonal entries.

Finally, for each input matrix (SC, FC, or CMs), AMI and MMS were computed across increasingly finer partitions by systematically varying the resolution parameter *γ* (Fig 1d). To account for the inherent stochasticity of the Louvain algorithm, each partition was computed 100 times, resulting in a distribution of AMI and MMS estimates for each value of *γ*. We operationalize resolution-resolved partition agreement as the peak and area-under-the-curve (AUC) of the similarity measures. The peak similarity is the optimal agreement between reference and empirical partition across all values of *γ*, while the AUC similarity summarizes multi-resolution partition agreement into a single value.

### Network communication models narrow the gap between structural and functional modules

Figure 2a shows the AMI and number of modules obtained for SC, FC, and 4 CMs, across a range of partition resolution parameters. We first observed that, in comparison to SC and CMs, FC led to markedly greater agreement to reference functional systems. This is expected as both partitions are derived from functional MRI and it corroborates previous studies on the modularity of functional brain networks [20]. We used the spin null model [58] to create a set of 50000 spatially rotated reference partitions and computed a surrogate distribution of similarities between original and rotated reference partitions. Apart from exceedingly small values of *γ*, empirical modules significantly outperformed the surrogates (non-parametric *p*-value *<* 0.05), indicating that the AMI observed for all input matrices cannot be trivially explained by spatial autocorrelation in functional cortical partitions [59, 60].

**FIG. 2.**
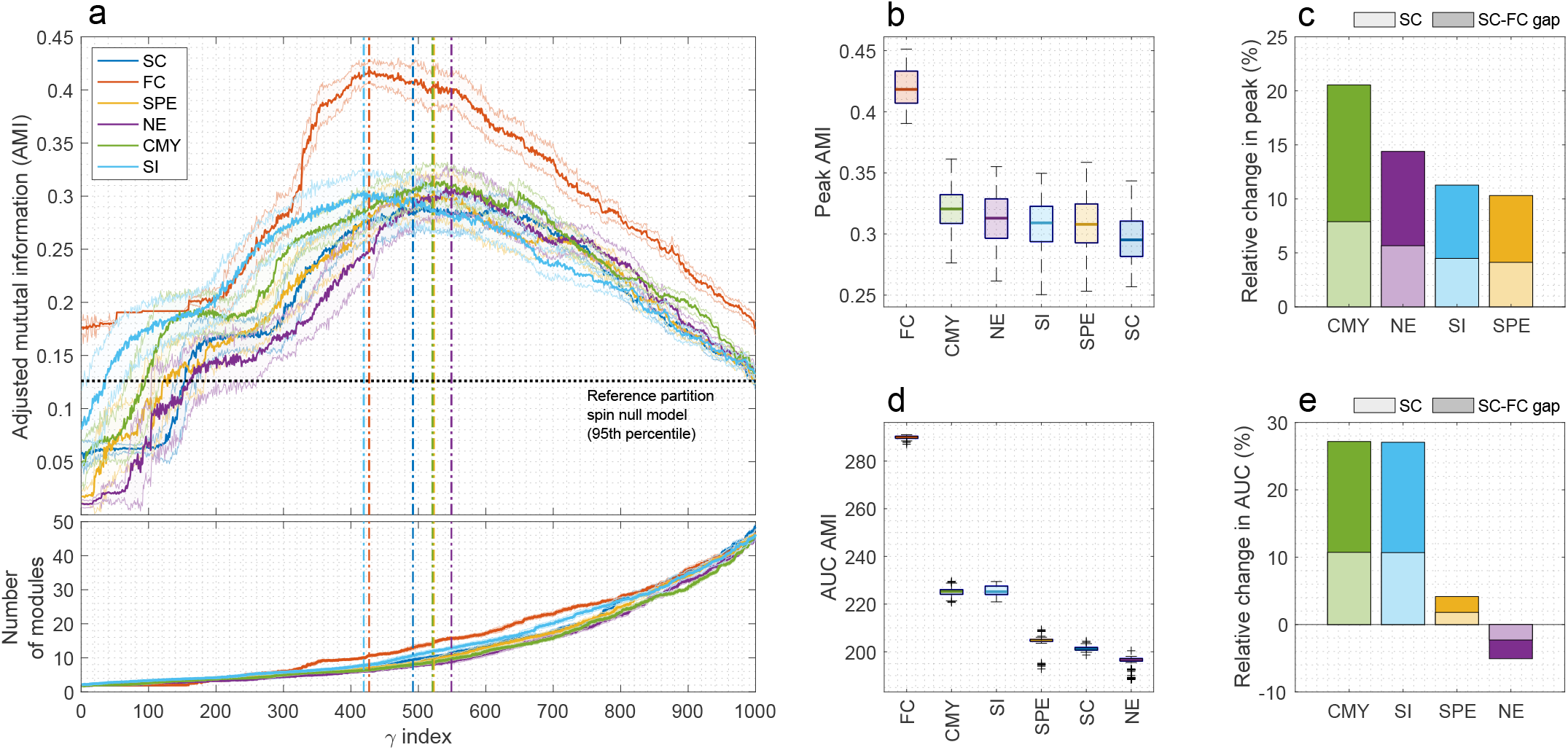
**(a)** Top: Adjusted mutual information (AMI) between empirical and reference partitions across a range of 1000 increasing resolution parameters *γ*. For each *γ* and for each input matrix, the average AMI (plus or minus one standard deviation) from 100 repetitions of the Louvain algorithm is shown. Horizontal dashed line indicates the 95% percentile of AMI expected under the null condition achieved by randomly rotating the reference partition. Vertical dashed lines mark the peak AMI for each input matrix. Bottom: average (plus or minus one standard deviation) number of modules obtained for each *γ*. Partitions identified by the Louvain algorithm become finer-grained as *γ* increases. We note that the horizontal axis refers to the *γ* index *k* = 1, …, 1000 of the resolution parameter sample yielding partitions with 2 to *N/*2 modules. The portion of the parameter space where this occurs is different for each input matrix, and thus the same *γ* index maps onto different *γ* values for SC, FC and CMs. **(b)** Peak AMI. **(c)** Light-colored bars show the relative change in peak AMI between SC and CMs. FC and SC were used as upper and lower benchmarks, respectively, of how well an empirical partition aligns with the reference. Dark-colored bars show what percentage of the SC–FC peak AMI gap is closed by CMs. **(d-e)** Same as (b-c) for the area-under-the-curve (AUC) AMI.

We found that all communication models outperformed SC when considering the best partitions obtained via systematically sweeping over the *γ* parameter space (Fig 2b; peak AMI two-sample t-test *p*-values *<* 10^−4^ [*df* =99] between SC and each model). The communicability model resulted in the CM with the modular decomposition most well aligned with functional modules, leading to an increase of 7.9% in peak AMI relative to SC (*p*-value *<* 10^−16^; Fig 2c, light-colored bars). To contextualize this improvement, we considered the peak AMI obtained from FC and SC as upper and lower benchmarks, respectively, for the correspondence to the reference. The increase in peak AMI afforded by the communicability model led to a narrowing of 20.5% of the gap between SC and FC performances (Fig 2c, dark-colored bars). Summarizing multi-resolution AMI estimates using the AUC (Fig 2d,e) resulted in top performances by the communicability and search information models, both improving on SC by approximately 11% (both *p*-values *<* 10^−16^) and narrowing the SC–FC gap by approximately 27%. Navigation was the only model that led to a significant decrease in SC’s AUC similarity to functional modules (−2.3%; *p*-value *<* 10^−8^).

Assessing partition agreement using MMS yielded comparable findings (Fig 3). As expected, FC resulted in the modular decompositions most well aligned with functional reference systems, while all empirical partitions significantly outperformed the spin null model for most of the *γ* parameter space (Fig 3a). Communicability remained the best performing communication model, contributing to significant improvements relative to SC for both peak and AUC MMS (5.1% and 7.8%, respectively; both *p*-values *<* 10^−8^; Fig 3c,e, light-colored bars). Once again, these improvements represented marked reductions of the SC–FC performance gap for both peak and AUC MMS (25.3% and 47.8%, respectively; Fig 3c,e, dark-colored bars).

**FIG. 3.**
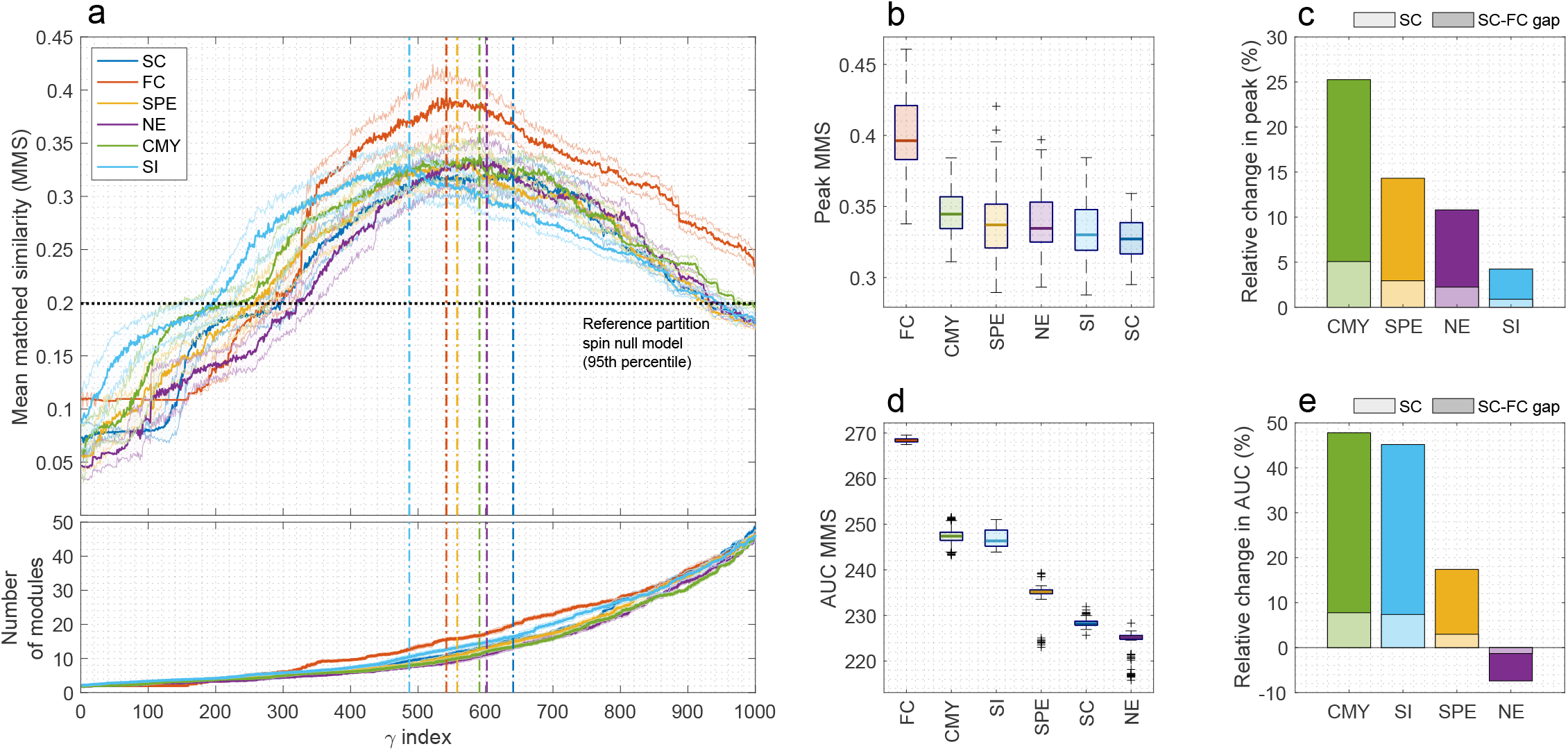
**(a)** Top: Mean matched similarity (MMS) between empirical and reference partitions across a range of 1000 increasing resolution parameters *γ*. For each *γ* and for each input matrix, the average MMS (plus or minus one standard deviation) from 100 repetitions of the Louvain algorithm is shown. Horizontal dashed line indicates the 95% percentile of MMS expected under the null condition achieved by randomly rotating the reference partition. Vertical dashed lines mark the peak MMS for each input matrix. Bottom: average (plus or minus one standard deviation) number of modules obtained for each *γ*. Partitions identified by the Louvain algorithm become finer-grained as *γ* increases. **(b)** Peak MMS. **(c)** Light-colored bars show the relative change in peak MMS between SC and CMs. FC and SC were used as upper and lower benchmarks, respectively, of how well an empirical partition aligns with the reference. Dark-colored bars show what percentage of the SC–FC peak MMS gap is closed by CMs. **(d-e)** Same as (b-c) for the area-under-the-curve (AUC) MMS.

These results indicate that modules derived from CMs, in particular for communicability, are more aligned with canonical functional systems than modules derived from SC. This was the case for both AMI (agnostic about module matching) and MMS (explicit about module matching) measures, as well as for peak (best *γ* for each input matrix) and AUC (summary across *γ* range) operationalizations of resolution-resolved partition similarity.

To further investigate these findings, we tested whether any partition resolutions could be found for which network communication models were detrimental to the agreement with reference functional systems. To this end, we computed two-sample *t*-tests comparing the performances of SC and CMs at each *γ* in the resolution parameter space (Fig 4). The resulting curves of *t*-statistics revealed that the impact of modeling network communication was not constant across resolutions, with peaks and troughs in model performance suggesting a complex, multi-scale interplay between the brain’s SC, communication dynamics, and mesoscale functional organization [61]. In fact, despite their overall benefit, most models contributed to decreasing (negative *t*-statistics) the agreement between SC and reference modules in at least some position of the parameter space. Crucially however, communicability was the only model that consistently contributed to explaining reference functional systems, with positive *t*-statistics across all resolutions of modular decomposition (with exceptions of marginal and brief negative values for *γ* indexes around 5 and 750). This was the case even when considering the resolution that optimized the similarity between SC modules and the reference (dashed dark blue lines in Fig 4; AMI: *t* = 5.4, *γ* index = 492, Fig 4; MMS: *t* = 0.04, *γ* index = 641, Fig 4).

**FIG. 4.**
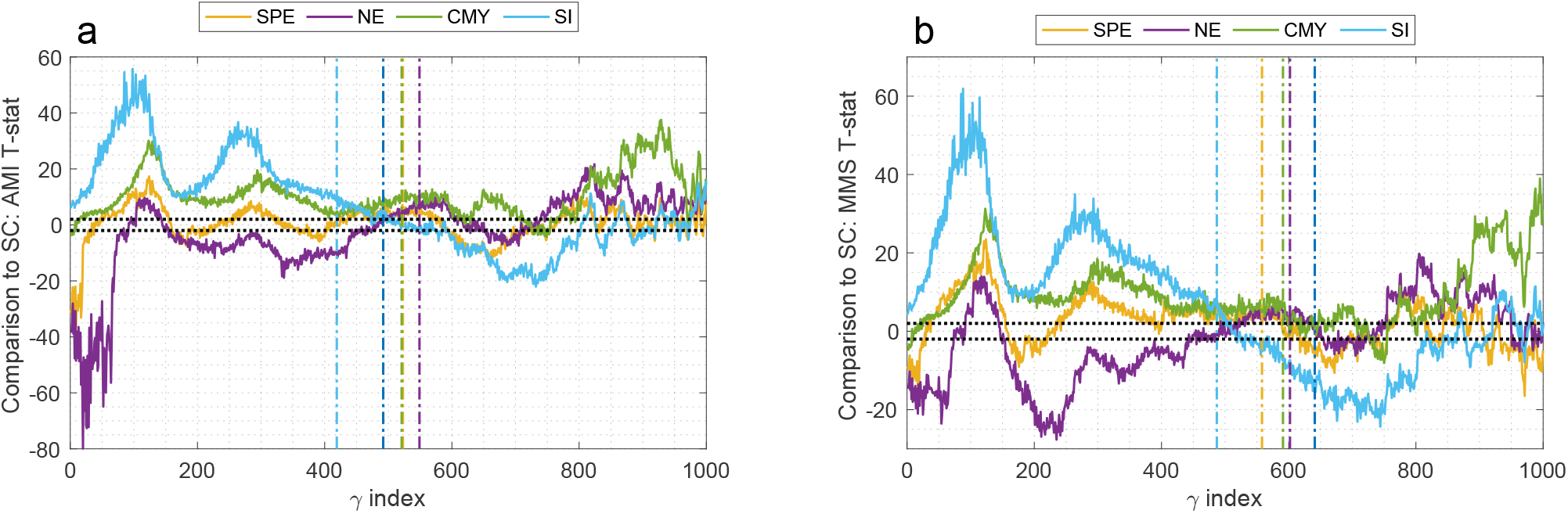
Statistical comparison between modules obtained for SC and CMs considering the **(a)** AMI and **(b)** MMS similarity measures. For each *γ*, a two-sample t-test was performed to compare the SC and CM distributions of partition similarity obtained via 100 repetition of the Louvain algorithm. Positive (negative) *t*-statistics indicate that modeling network communication increased (decreased) the similarity between structural and functional modules. Horizontal dotted black lines mark |*t*| = 2, denoting statistically significant differences at *α* = 5%. Vertical dashed lines mark the peak (a) AMI and (b) MMS for each input matrix.

Collectively, our results indicate that modeling polysynaptic communication on top of SC can help further explain the emergence of canonical functional systems from the underlying substrate of anatomical connectivity. In general, the benefits afforded by communication models were more pronounced when synthesizing results across community decomposition scales than when considering specific resolutions of modular organization. Critically, communicability—the top performing model across all scenarios explored—augmented SC’s account of canonical functional systems across the entire resolution parameter space, suggesting that this communication model captures the formation of functional components at multiple scales of investigation.

### Impact of network communication models on the identification of individual functional systems

We investigated the extent to which modules obtained for SC and CMs recapitulated specific functional systems of the reference partition. Under our module matching framework (Fig 1c), diagonal entries of the matched similarity matrix quantify the agreement between individual reference and empirical modules.

Figure 5a shows the AUC of the matched similarity obtained for each reference functional system. We observed considerable variation in the performance of communication models depending on the functional system. Modeling communication was most beneficial to the identification of the somatomotor, limbic and default mode systems, for which all CMs outperformed SC. In contrast, all communication models were detrimental to the characterization of the attention ventral network. For the visual, attention dorsal, and control systems, results were dependent on the choice of communication model, with communicability once again featuring as the model yielding the most robust benefits (Fig 5b). More specifically, communicability modules improved the match to somatomotor (20.3% increase in matched similarity relative to SC), limbic (+17.4%), attention dorsal (+7.0%), visual (+6.5%), default mode (+5.2%) and frontoparietal control (+1.2%) systems, while decreasing the match to the attention ventral system (−17.8%).

**FIG. 5.**
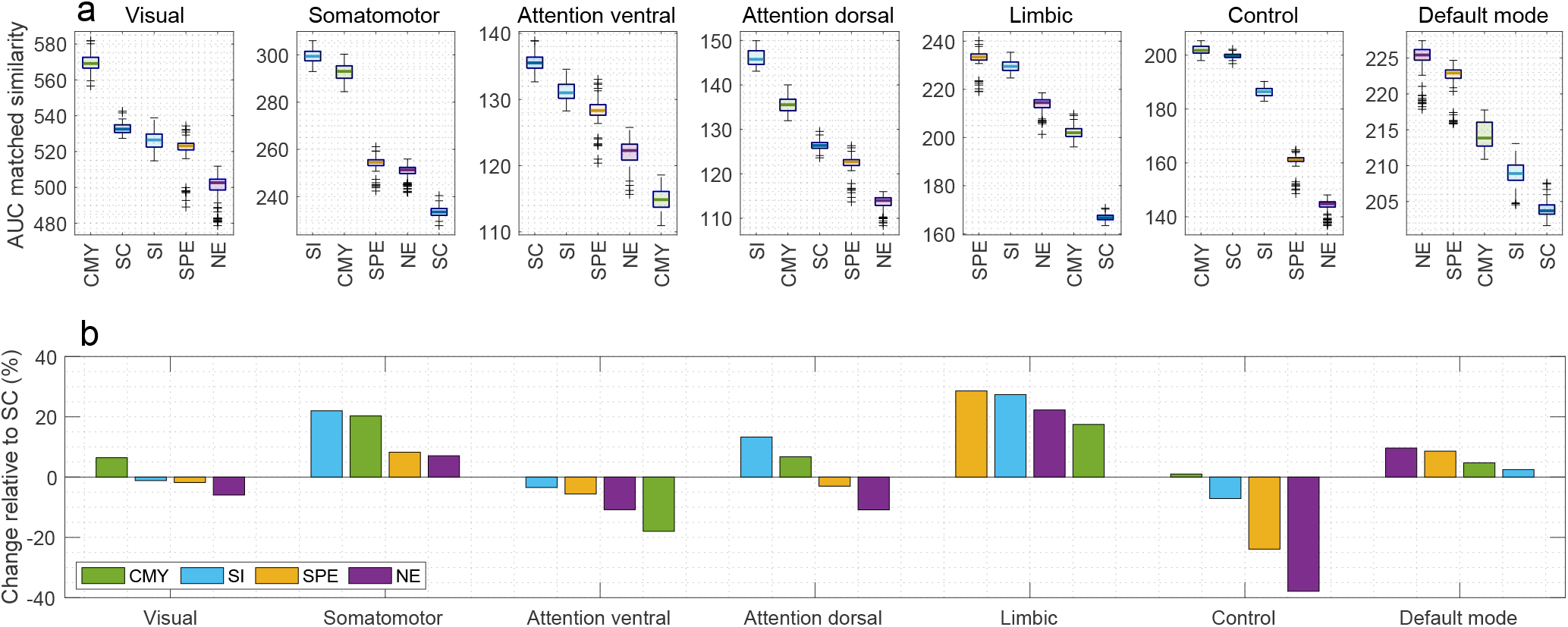
Impact of network communication models on the identification of individual functional systems. **(a)** AUC matched similarity obtained for the 7 functional modules of the ground truth. **(b)** Relative change in AUC matched similarity between SC and CMs.

To gain further insight into these results, we next sought to visualize SC and CM partitions. Here, it is important to reiterate that the benefits of modeling communication were more pronounced when considering the AUC across resolution parameters than for any individual *γ*. Visualizing the improved alignment to functional systems afforded by CMs is therefore challenging, since visualizing partitions requires the choice of a single modular scale. In addition, as seen in Fig 4, the choice of *γ* influences the extent to which CMs increase or decrease the match to the functional reference. Having noted these caveats, we investigated exemplar SC and communicability partitions obtained for a representative *γ* index 492 (corresponding *γ* values of 0.3484 and 0.5286, for SC and communicability, respectively). This resolution maximized the AMI between the SC and reference partitions (Figs 2a, 4a), and was chosen to provide a parsimonious comparison that does not inflate the benefits of modeling network communication.

Figure 6a,b illustrates the module matching process that assigned SC and communicability modules to canonical functional systems. Since the matching is strictly one-to-one and the reference partition contains 7 functional systems, 3 out of 10 SC modules and 2 out of 9 communicability modules remained unmatched. Following module matching, the communicability partition better recapitulated the somatomotor (SM; 31.2% increase in similarity), attention dorsal (AD; +35.4%), limbic (LIM; +16.7%), control (CON; +58.3%) and default mode (DMN; +70.0%) systems, while SC modules provided a better fit for the visual (VIS; -17.6%) and attention ventral (AV; -12.5%) systems (Fig 6c).

**FIG. 6.**
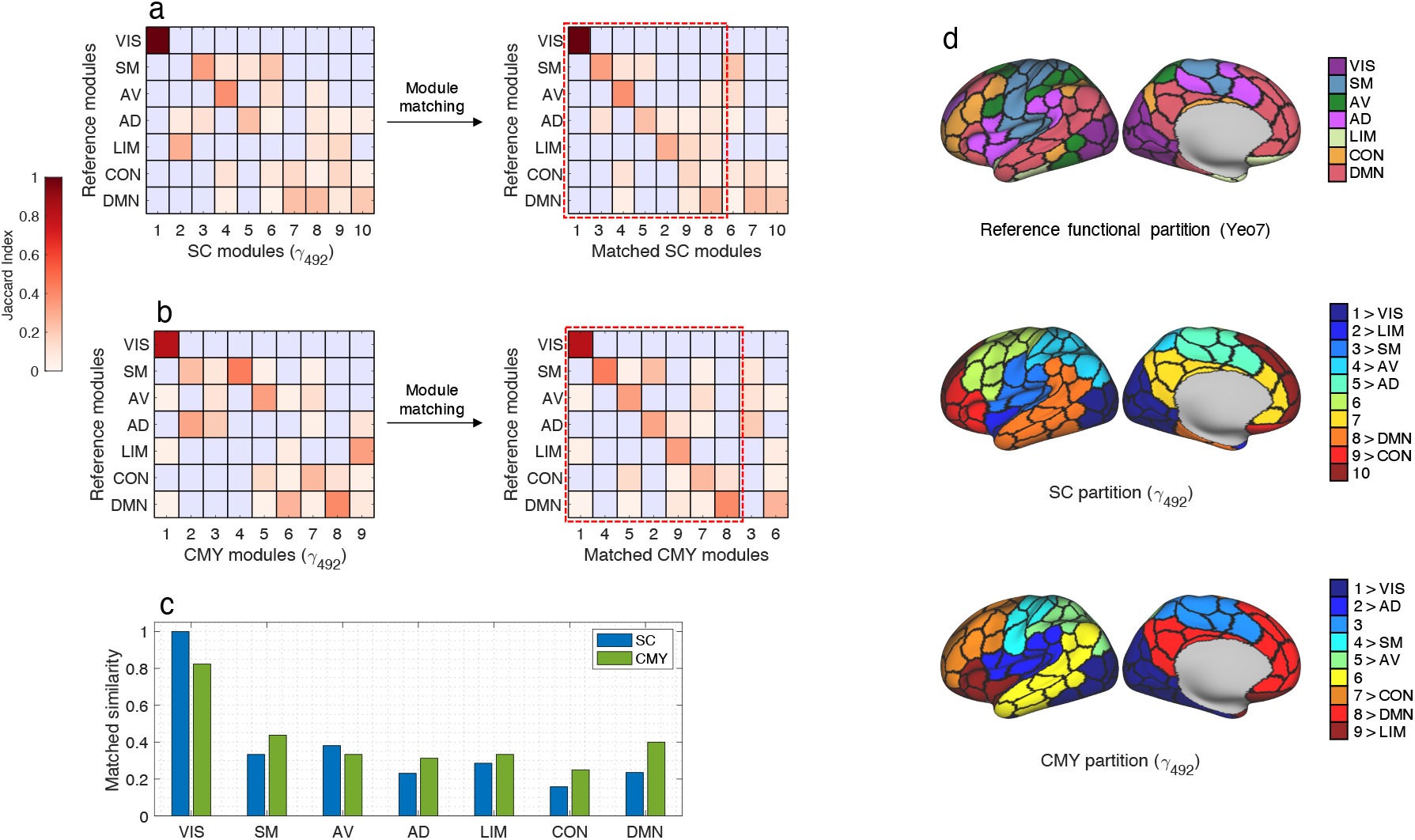
Comparison of SC and communicability partitions obtained for *γ* index 492 (corresponding to the peak AMI between the SC and reference partitions). Similarity and matched similarity matrices for the **(a)** SC and **(b)** communicability partitions. Each matrix entry is the Jaccard index between a pair of reference (rows) and SC (columns) modules. A one-to-one matching between reference and empirical modules is performed by reordering the columns of the similarity matrix with the goal of maximizing the sum of the main matrix diagonal. The red dashed square delineated the 7-by-7 matrix of matched modules, while empirical modules outside the square remained unmatched. **(c)** Diagonal entries of the matched similarity matrices representing the alignment to individual reference functional systems. **(d)** Cortical maps of the reference, SC and communicability partitions. The colour keys show the pairings obtained from the module matching routine.

Figure 6d shows the cortical maps obtained by projecting the functional reference, SC and communicability partitions onto the cortical surface. We first observed that, as with SC modules, the communicability partition remained contiguous in space, unable to unify distant parts of functional systems into spatially distributed modules. For example, the medial (including precuneus and posterior cingulate cortex) and temporal-parietal components of the DMN were well characterized by the communicability partition, but they were assigned to separate modules (CMY modules 8 and 6, respectively). As such, instead of large-scale rearrangements, communicability promoted local adjustments to the modular organization of SC. While spatially localized, these changes were consequential, leading to a communicability partition that better recapitulated the individual canonical systems of the reference (Fig 6c). For instance, regions in the central sulcus area were assigned to 3 adjacent modules in the SC partition (SC modules 3, 4 and 6). The polysynaptic interactions estimated by the communicability model reshaped the local modular organization around the central sulcus, reassigning certain regions of the adjacent SC modules into a new community (CMY module 4) that more closely matched the boundaries of the somatomotor system.

### Control analyses

We performed control analyses to test whether the key findings reported in the main text were robust to a range of methodological choices. Specifically, we considered variations in: (i) cortical parcellation (Schaefer *N* = 200 [main] vs. Glasser *N* = 360 [control]); (ii) pipeline of fMRI time series pre-processing (global signal regression [main] vs. no global signal regression [control]); (iii) null models of modularity maximization for SC (Potts [main] vs. Girvan–Newman [control]); and (iv) the definition of the AUC similarity measures (computed across the entire resolution parameter space [main] vs. computed across the range of *γ* values for which empirical partitions outperformed the spin null model of spatially rotated reference partitions [control]).

For each of these scenarios, we recomputed the entire multi-scale community detection routine for SC, FC and CM input matrices. Following the analyses performed in Figures 2 and 3, we derived the peak and AUC of the AMI and MMS to the reference partition. Figures S2 (AMI) and S3 (MMS) summarize the impact of modeling network communication in relation to the empirical partitions obtained for SC and FC. Our findings were consistent across all control analyses. Modeling communicability on top of SC reliably improved the alignment to reference functional systems (4–15% increased similarity) and narrowed the gap between SC and FC empirical partitions (6–60% reduction of the gap). The only exception was the peak MMS for the Glasser parcellation, for which no significant difference between communicability and SC was observed. We also note that for a number of scenarios the search information model performed comparably to communicability.

## DISCUSSION

We provide multiple lines of evidence suggesting that network communication models can help explain the emergence of canonical functional systems from anatomical connectivity. Structural connectivity (SC) matrices inferred from tractography encode direct anatomical connections between pairs of regions and are typically sparse, given that most regional pairs are not directly connected. Pairs of regions that are not directly connected must communicate polysynaptically—via one or more intermediary regions. A network communication model determines a strategy to guide polysynaptic signal propagation. Under a given model, the ease of communication between each pair of regions can be encoded in the communication matrix (CM), which is derived from SC using analytical transformations. Whereas SC is typically sparse, the CM is dense and reveals information about all pairs of regions, not just those that are directly connected.

A growing body of evidence supports that CMs can explain a greater portion of variation in FC than SC alone. Previous studies along these lines have focussed mostly on the global scale [30, 31]—FC between regions pairs across the whole brain—and local scale [62, 63]—FC profiles of individual regions. Here, we add to this literature by considering how network communication models can uncover mesoscopic relationships between brain structure and function.

We found that, in comparison to SC, the modularical connectivity. Structural connectivity (SC) matrices inferred from tractography encode direct anatomical connections between pairs of regions and are typically sparse, given that most regional pairs are not directly connected. Pairs of regions that are not directly connected must communicate polysynaptically—via one or more intermediary regions. A network communication model determines a strategy to guide polysynaptic signal propagation. Under a given model, the ease of communication between each pair of regions can be encoded in the communication matrix (CM), which is derived from SC using analytical transformations. Whereas SC is typically sparse, the CM is dense and reveals information about all pairs of regions, not just those that are directly connected.

A growing body of evidence supports that CMs can explain a greater portion of variation in FC than SC alone. Previous studies along these lines have focussed mostly on the global scale [30, 31]—FC between regions pairs across the whole brain—and local scale [62, 63]—FC profiles of individual regions. Here, we add to this literature by considering how network communication models can uncover mesoscopic relationships between brain structure and function.

We found that, in comparison to SC, the modular structure of CMs more closely recapitulates the mesoscale functional organization of the brain into intrinsic restingstate networks. Therefore, by taking into account polysynaptic communication, we approximate structural modules—primarily influenced by neuroanatomical proximity [7]—to functional modules—reflecting patterns of interregional co-activity implicated in specific cognitive domains [24]. Our results provide insight into how the interplay between anatomical wiring, neural communication dynamics, and the emergence of specialized components of information processing.

Importantly, which brain network communication models most accurately and parsimoniously describe biological signaling remains an open question [28]. We considered four previously explored candidate models, chosen to cover a range of putative neural signaling strategies. Shortest path and navigation are routing models that propose where communication between two regions takes place via a single, highly efficient path. While advantageous from the perspectives of transmission delays and metabolic costs, efficient routing hinges on the assumption that individual nodes have knowledge about the network beyond their immediate vicinity, a requirement that might not be realistic for decentralized systems such as the brain [64]. In contrast, search information and communicability stem from a diffusive conceptualization of network communication. Signal propagation through diffusion does presuppose the same knowledge assumptions, but it typically entails longer transmission delays and higher energy expenditure [65].

Of the communication models investigated, communicability resulted in partitions with the highest correspondence to functional canonical systems—a finding that was observed across most resolutions of modular decomposition and individual reference systems (with the exception of the ventral attention network). Critically, partition of the communicability CM yielded modules that better recapitulated canonical functional systems than partitions identified in the SC matrix. Taking into account results obtained from different similarity measures, cortical parcellations, fMRI preprocessing pipelines, and modularity maximization models, we found that communicability generally accounts for 20 to 60% of the gap between SC and FC partitions.

Communicability is a diffusion-based model whereby information is “broadcast” along all possible sets of connections that link two regions [38, 66], contrasting with other models predicated on a single communication path that is selectively accessed. This type of diffusive communication may be better suited to integrate information between near-by regions, since broadcast strength diminishes as a function of topological distance [2]. This aligns with the observation that communicability’s increased match to functional systems was the result of spatially localized adjustments to the SC’s modular structure, instead of the unification of distant regions into spatially distributed modules. Therefore, we conjecture that the benefit of diffusive models stems from a more comprehensive utilization of local network topology—instead of utilization of direct connections alone or single efficient paths—in the communication between regions in close topological proximity, which in turn promoted functionally meaningful refinements of modular boundaries.

Our findings are in close agreement with previous work investigating systems-level brain network communication. Betzel and colleagues reported that detecting communities using a diffusion random walk model accounted for unimodal functional systems such as the visual and somatomotor networks [61]. Using a graph matching approach, Osmanlioglu and colleagues found that communicability, in comparison to other communication models, provided the most accurate description of systems-level FC [67]. We extend these efforts through a direct comparison of structural, functional and communication partitions identified using modularity maximization. More broadly, our work also intersects with theoretical and computational research on the use of random walks to identify communities in complex networks [68].

Finally, our results corroborate previous reports on the utility of communicability to investigate a range of diverse neuroscience questions [39]. Examples include studies on the impact of stroke lesions [69], effects of neurodegeneration [70], simulations of neural gain fluctuations [71], and pharmacogenetic manipulation of brain regions [72]. More generally, we add to mounting empirical evidence challenging the notion that communication in brain networks occurs exclusively via topological shortest paths [30, 31, 73], an assumption built into many popular graph measures in network neuroscience (e.g., betweenness centrality or global efficiency).

### Technical considerations and alternative approaches to link structural and functional modules

SC is sparse while CMs and FC are fully connected. Two different modularity maximization null models are typically used in these cases—the standard Girvan-Newman model for sparse, and the Potts model for dense matrices [9]. To ensure that changes in partition similarity did not result from different definitions of modularity maximization [74], we repeated our analyses using both null models to identify communities in SC. Relatedly, we note that it is unlikely that the communicability CM improves on SC trivially due to its higher connection density, since, while all CMs are full graphs, only communicability led to consistent benefits.

While we focussed on network communication, a number of alternative approaches have been used to investigate the relationship between structural and functional modules. Modularity maximization null models that correct for wiring cost have been shown to yield structural partitions that accurately recapitulate spatially distributed functional systems, such as the frontoparietal control network [75]. In a recent paper, Puxeddu and colleagues introduced a multi-layer community detection framework that facilitates the comparison of partitions obtained from different input networks [76]. Using this approach, the authors found that the relationship between SC and FC partitions is influenced by the resolution in which modular organization is investigated.

Other examples include the use of biophysical models of neural dynamics [47], hierarchical clustering [77], multivariate statistical techniques [17], and block models capable of identifying non-assortative modular structures [78, 79]. Our work complements these efforts from the perspective of network communication. The CMs explored here can be readily integrated to advanced statistical and community detection methods to investigate synergies between these parallel lines of research.

### Limitations and future directions

Following previous work on brain network modeling [42, 43], our analyses were performed on intra-hemispheric networks. Poor reconstruction of cross-hemispheric fibers is a well-documented issue in tractography [44] with important implications for structure-function analyses [80, 81]. We also note that canonical functional systems are spatially distributed both within as well as homotopically between hemispheres. In the absence of reliable cross-hemispheric SC, restricting our analyses to single hemispheres mitigated the possibility of confounding intra- and inter-hemispheric partition similarity. More broadly, variations in SC mapping methods can impact the computation of network communication models [82–84] and future work is necessary to replicate our findings in alternative reconstructions of structural brain networks.

We adopted the 7 resting-state networks defined by Yeo and colleagues as the reference partition for the mesoscale functional organization of the human brain [22]. In the last decade, this partition has become firmly established as a functional taxonomy of cortical regions [45–51]. Nonetheless, the choice of this—or any alternative [20]—functional reference is inevitably an oversimplification, as brain function is context-dependent, subject-specific, and unfolds at multiple spatial and temporal scales. Understanding how structure mediates the emergence of functional components in a wider range of scenarios is an important direction for future work.

A limitation of our one-to-one matching approach is that it penalizes cases where a partition accurately clusters together the regions of a functional system but splits them into one or more modules (see Fig 6). Partition similarity measures is an active topic of technical research [56] and developments in this area could provide more suitable alternatives to the quantification of the match between structural and functional modules.

## Conclusion

In summary, we demonstrated that taking into account polysynaptic interactions via models of network communication can narrow the gap between the mesoscopic organization of structural and functional connectivity. Our work provides new insights into systems-level properties of brain networks and contributes to the understanding of large-scale neural communication.

## MATERIALS AND METHODS

### Brain connectivity mapping

#### Structural connectivity

Minimally preprocessed high-resolution diffusion-weighted MRI from the HCP were used to map structural brain networks for 1000 healthy young adults. Acquisition and preprocessing details of diffusion MRI data are described in [85, 86]. For each participant, whole-brain white matter tractograms were mapped using a probabilistic tractography pipeline implemented in MRtrix3 [87] (multi-shell multi-tissue constrained spherical deconvolution [88], iFOD2 tracking algorithm [89], anatomically constrained tractography [90], 5M streamlines; further details in [91]). Cortical gray matter regions were delineated according to the Schaefer (*N* = 200; main text) [41] and Glasser (*N* = 360; control analysis) [92] parcellations. The connection weight between a pair of gray matter regions was defined as the number of streamlines connecting them divided by the product of their surface areas. Connections with fewer than 5 streamlines were discarded to attenuate the high false positive rate of probabilistic tractography [84]. Individual structural networks were combined into a group-consensus SC matrix that preserved the average connection density across participants (41% and 21% for the Schaefer and Glasser parcellations, respectively) [93].

#### Functional connectivity

Minimally preprocessed ICA-FIX resting-state functional MRI data for the same 1000 participants were acquired from the HCP. For each participant, four (2 sessions on separate days with both right-to-left and left-to-right phase encodings) 14 minutes and 33 second scans (0.72s TR) were collected. Details on resting-state protocol and preprocessing are provided in [85, 94]. FC was computed according to two different pipelines. In the first pipeline (main text), voxel-level blood-oxygen-level-dependent (BOLD) time series were linearly detrended, band-pass filtered, and standardized [95]. Next, four nuisance variables were regressed out: the global signal (GS), the GS squared, the GS derivative, and the squared GS derivative [96]. In the second pipeline (control analysis), no further preprocessing was performed to the minimally preprocessed ICA-FIX data from the HCP. In both pipelines, the time series of voxels within the same gray matter region were averaged and FC was computed as the Pearson correlation between regional time series. Group-level FC matrices were computed by averaging a total of 4,000 matrices (4 per subject for 1000 subjects) for the Schaefer and Glasser parcellations.

### Network communication models

Network communication models were computed using the Brain Connectivity Toolbox [97]. Computations were carried out separately for left and right intra-hemispheric networks. Let *W* ∈ ℝ^*N×N*^ be the matrix of structural connectivity weights between *N* regions, where *W*_*ij*_ = 0 and *W*_*ij*_ *>* 0 denote, respectively, the absence and presence of a connection between regions *i* and *j*. We define a matrix of structural connectivity lengths *L* =− log_10_(*W*) [98]. While W measures the strength and reliability of anatomical connections supporting communication, L quantifies the distance or travel cost between regions [30]. The transformation of connection weights into lengths is necessary to the computation of network communication models that seek to minimize the cost of communication between regions.

#### Shortest path efficiency

Let Λ* denote the matrix of shortest path lengths, where 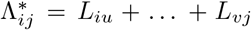 is the sum of connection lengths traversed along the shortest path between regions *i* and *j*. The shortest path efficiency CM was defined as *SPE* = 1*/*Λ* [34].

#### Navigation efficiency

Navigation implements a greedy routing strategy based on a measure of nodal distance. Following previous work, we used the Euclidean distance between region centroids to identify navigation paths [36]. Starting from a source region *i*, navigation progresses to *i*’s neighbor that is closest in distance to a target region *j*. This simple rule is repeated until *j* is reached (successful navigation) or a region is revisited (failed navigation). Successful navigation path lengths are defined as Λ_*ij*_ = *L*_*iu*_+…+*L*_*vj*_, i.e., the sum of connection lengths traversed along the navigation path from *i* to *j*, whereas failed navigation yields Λ_*ij*_ = ∞. The navigation efficiency CM was defined as *NE* = 1*/*Λ.

#### Search information

Search information measures the accessibility of efficient communication paths in a network [37]. It is computed based on the probability of a random walker serendipitously traveling between two regions via their shortest path. An unbiased random walker travels from region *p* to region *q* with probability *T*_*pq*_ = *W*_*pq*_*/s*_*p*_, where *s*_*p*_ is the (outgoing) strength of *i*. The probability of the random walker travelling from *i* to *j* along their shortest path {*i, u*, …, *v, j*} is given by *P*_*ij*_ = *T*_*iu*_ ×…×*T*_*vj*_. Search information is typically defined as *SI*_*ij*_ = −*log*_2_(*P*_*ij*_) [31], with higher values of *SI*_*ij*_ indicating that efficient routes from *i* to *j* are less accessible. However, community detection requires matrices that encode the propensity of region pairs to belong to the same module. We therefore defined the search information CM as *SI′*= 1*/SI*.

#### Communicability

Communicability is defined as a weighted sum of the total number of walks between two nodes [38]. Formally, the 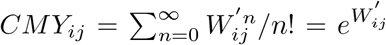. Weighted connectivity matrices are typically normalized prior to the computation of communicability to attenuate the influence of high strength nodes, such that 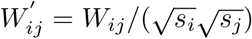 and *s*_*i*_ is the strength of node *i* [99].

### Multi-resolution community detection

#### Modularity maximization

or a given resolution parameter *γ* and input matrix *A*, community detection was computed as follows. First, *A* was symmetrized (certain CMs can be asymmetric [32]), z-scored, and shifted by the absolute magnitude of the minimum z-scored entry, thus ensuring that all matrix entries are non-negative. These steps avoid the need for special modularity maximization null models to handle the negative connection weights of FC and contribute towards a comparable range of *γ* values across input matrices. The Brain Connectivity Toolbox implementation of the Louvain algorithm was then used to maximize the modularity statistic

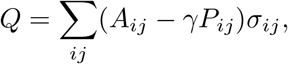

where *P*_*ij*_ is the expected weight of the *ij* connection under a null model *P*, and *σ*_*ij*_ = 1 if *i* and *j* are assigned to the same module and *σ*_*ij*_ = 0 otherwise.

The choice of *P* is typically influenced by the properties of *A*. Previous work indicates that the Potts (also known as uniform) model [100] *P* = **1**^*N×N*^ is suitable for dense matrices, such as FC and CMs, in which connection weights are not independent from each other [101]. On the other hand, sparse matrices such as SC are typically clustered using the Girvan–Newman (also known as configuration) model 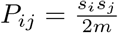, where *s*_*i*_ is the strength of node *i*, and *m* is the total number of connections in the network [102, 103]. The definition of *P* also has implications to the interpretation of modules uncovered with modularity maximization [9, 74]. Under the Girvan–Newman model, two nodes tend to be assigned to the same module if they are more strongly connected than expected based on their combined connectivity to the rest of the network. Meanwhile, the Potts model presupposes that every node pair is connected with the same weight, leading to partitions in which within-module connectivity is, on average, larger than *γ*. In order to compare the modular organization of SC, FC and CMs on an equal footing, in the main text, we used the Potts model to perform community detection for all input matrices. For completeness, we performed control analyses in which we used the Girvan–Newman model to cluster SC.

Two-pass multi-resolution routine. Following previous work [52–54], the multi-resolution community detection for an input matrix *A* was implemented as follows. In the first pass of the routine, we sampled 100 values of the resolution parameter 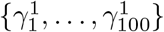, such that 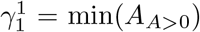 and 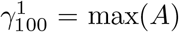. For each 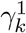, modularity maximization was performed using the Potts model as described above. We identified the 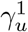 and 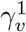 that resulted in, respectively, the first partition with at least 2 modules and the last partition with fewer than (*N/*2) + 1 modules.

In the second pass of the routine, we sampled 1000 values 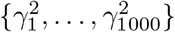, where 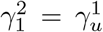 and 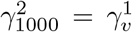. By construction, for each 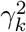, this resulted in partitions ranging from 2 to *N/*2 modules. In order to take into account the inherent stochasticity of the Louvain algorithm, we computed a set of 100 partitions for each 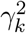. This routine was computed separately for left and right intra-hemispheric networks.

In both passes, values of *γ* were sampled linearly for FC and logarithmically for SC and the CMs, in order to provide good coverage of the weight distribution of each input matrix. Figure S1 shows a visualization of the *γ* sampling of the two-pass routine for each input matrix. The horizontal axes of Figures 2a, 3a and 4a,b refer to the index *k* of the *γ*^2^ sample. Note that the same *γ* index *k* maps onto different values of *γ* used for each input matrix.

### Partition similarity measures

The correspondence between the functional reference and an empirical partition was assessed by the adjusted mutual information and the mean matched similarity. Similarities were computed separately for left- and right-hemispheric partitions and averaged.

#### Adjusted mutual information

Consider two disjoint partitions of *N* regions *U* = {*U*_1_, …, *U*_*R*_} with *R* modules and *V* = {*V*_1_, …, *V*_*C*_} with *C* modules. The mutual information between U and V is defined as

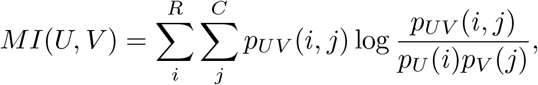

where *p*_*U*_ (*i*) = |*U*_*i*_|/*N* is the probability that a random region belongs to module *U*_*i*_ in partition *U*, and *p*_*UV*_ (*i, j*) = (|*U*_*i*_ ∩*V*_*j*_ |)*/N* is the probability that a random region belongs to module *U*_*i*_ in partition *U* and module *V*_*j*_ in partition *V*. The mutual information quantifies the similarity between U and V as the amount of information shared between the partitions. If *MI*(*U, V*) = 0, no information is shared between *U* and *V*, and thus knowledge of *U* provides no insight into *V* (and vice-versa). Meanwhile, if *MI*(*U, V*) = 1, knowledge of *U* provides complete insight into *V*. However, for two random *U* and *V, MI*(*U, V*) tends to grow as a function of *R* and *C*, thus overestimating the similarity between partitions with large numbers of modules.

The adjusted mutual information is a variation of the mutual information that corrects similarity solely due to chance [57]. It is defined as

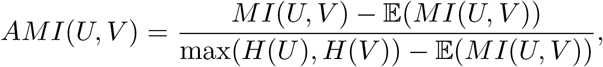

where 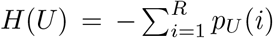 is the entropy of partition *U*, and 𝔼(*MI*(*U, V*)) is the expected value of the mutual information between *U* and *V* derived from a hypergeometric distribution (see [57] for further details). The AMI is 1 for identical *U* and *V* and 0 when the mutual information between them equals the value expected due to chance alone. The code used to compute AMI is available at [104].

#### One-to-one module matching

First, we computed the similarity matrix *S* ∈ ℝ^7*×n*^, where *S*_*ij*_ = |*U*_*i*_ ∩*V*_*j*_ |*/*|*U*_*i*_ ∪*V*_*j*_| is the Jaccard index between reference module *U*_*i*_ and empirical module *V*_*j*_, *i* ∈{1, …, 7 }, and *j* ∈{1, …, *n* }. The one-to-one matching of *U* and *V* modules was performed using the Munkres (Hungarian) algorithm for the linear assignment problem [105] applied to the dissimilarity matrix *D* = 1 −*S*. The algorithm identifies the reordering of rows (if *n <* 7) or columns (otherwise) of *D* that minimizes the sum of the main diagonal of the reordered dissimilarity matrix *D ′* We define the matched similarity matrix as *S′* = 1 −*D′* To summarize the similarity between *U* and *V* into a single value, we defined the mean matched similarity as 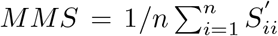 The code implementing the Munkres (Hungarian) algorithm is available at [106].

### Spin null model

Using an implementation of the spin test for cortical parcellations [59], we performed 50000 random spherical rotations of the reference functional partition. We computed the similarity between the original reference and the spatially rotated ones, yielding a null similarity distribution. An empirical partition was considered to outperform the spin null model if its similarity to the original reference exceeded the 95th percentile of null similarity distribution (non-parametric statistical significance test at *α* = 5%), marked by the horizontal dotted black lines in Figures 2 and 3. We note that this null model is particularly stringent, as surrogate partitions always have the same number of modules as the reference. Evidence that an empirical partition passes the spin test indicates that its similarity to the reference is not trivially explained by spatial autocorrelations inherent to cortical maps [58, 60].

## ACKNOWLEDGMENTS

We thank Joshua Faskowitz for valuable contributions to data preprocessing. Brain imaging data were provided by the Human Connectome Project, WU–Minn Consortium (1U54MH091657; Principal Investigators: David Van Essen and Kamil Ugurbil) funded by the 16 National Institutes of Health (NIH) institutes and centers that support the NIH Blueprint for Neuroscience Research; and by the McDonnell Center for Systems Neuroscience at Washington University. CS was supported by the Australian Research Council (grant number DP170101815); SML is supported by a Melbourne Research Scholarship; AZ is supported by the National Health and Medical Research Council of Australia (APP1118153); and FC is supported by the National Health and Medical Research Council of Australia (grant number APP1117724).

---

**FIG. S1.**
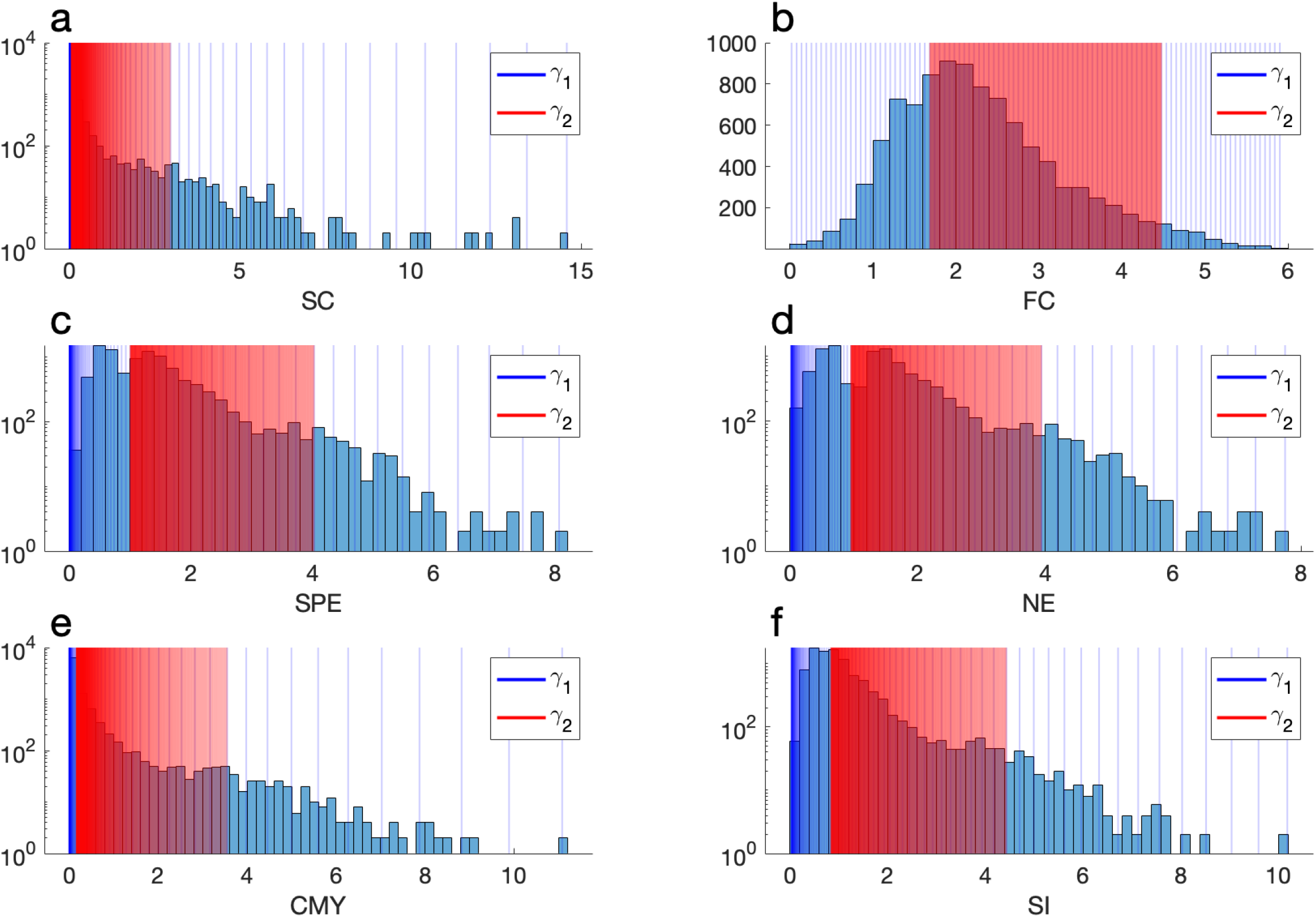
Samples of the *γ* parameter space used in the two-pass multi-resolution community detection routine. Histograms show the distribution of the z-scored input matrices. Matrices were z-scored to result in samples drawn across comparable ranges of parameter space. Blue lines mark the 100 parameter values 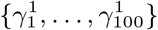 used in the first pass of the routine. We identified the 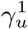 and 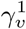 that resulted in, respectively, the first partition with at least 2 modules and the last partition with fewer than (*N/*2) + 1 modules. Red lines mark the 1000 parameter values 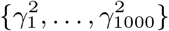, where 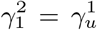 and 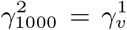. The resolution parameter space was sampled logarithmically to account for the skewed distribution of input matrices, with the exception of FC (panel b), for which *γ* values were sampled linearly.

**FIG. S2.**
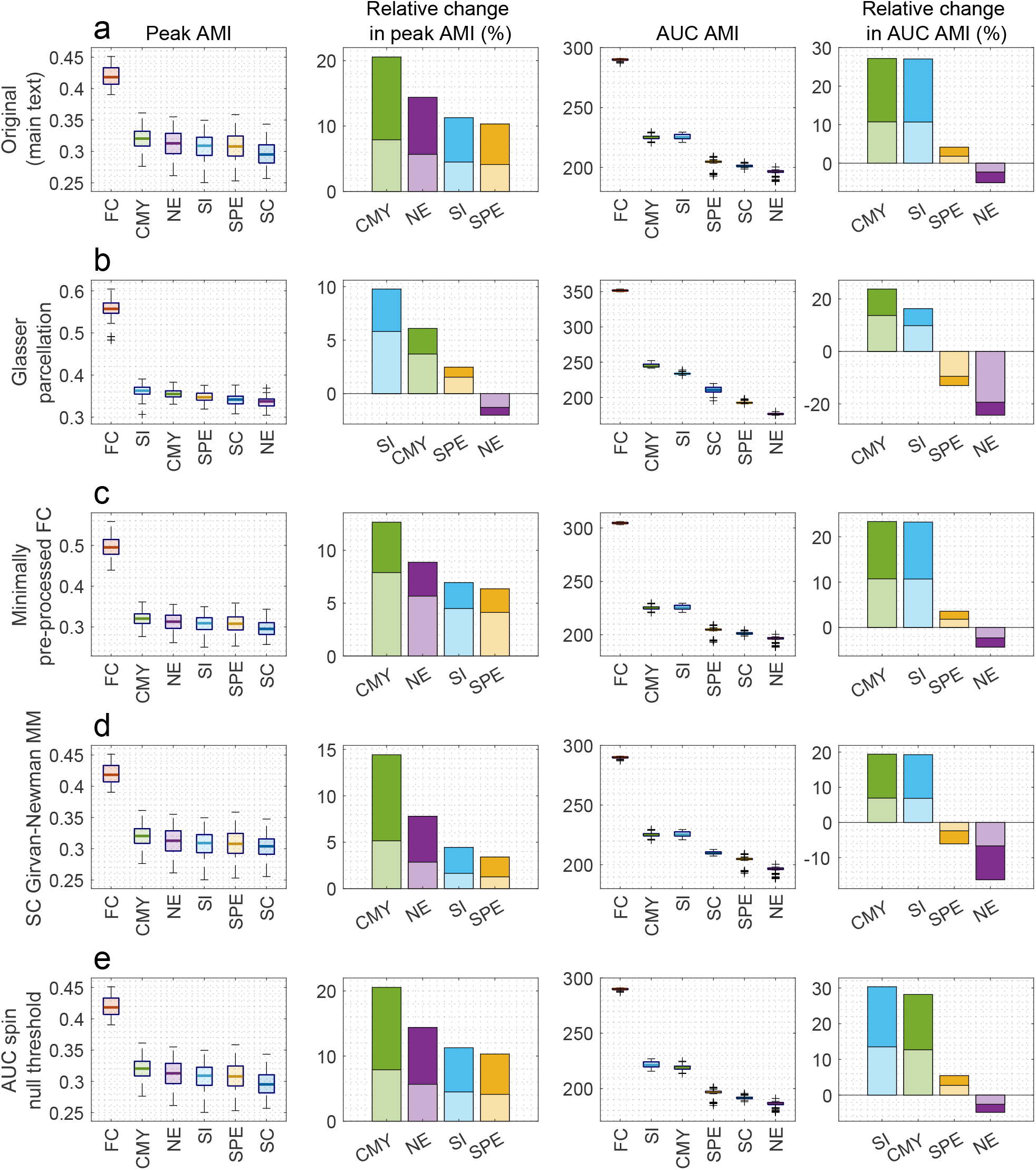
Control analyses using the AMI partition similarity measure. To facilitate the comparison of control analyses to the results presented in the main text, panel **(a)** recapitulates the results shown in Fig. 2b,c,d,e (respectively, left to right). The main text analysis was conducted considering (i) the Schaefer 200 parcellation, (ii) fMRI data pre-processed to regress out the mean global signal and other nuisance variables, (iii) SC matrices clustered using the Potts null model for modularity maximization, and (iv) the AUC summary of multi-resolution partition similarity computed across the entire resolution parameter space. In each control analysis, we tested the robustness of our results to changes in one of these four factors, while the other factors remained unaltered. **(b)** Control analysis using the Glasser parcellation. **(c)** Control analysis using the ICA FIX minimally preprocessed fMRI data from the Human Connectome Project, with no additional processing steps. **(d)** Control analysis using the Girvan–Newman null model for the modularity maximization of SC matrices. **(e)** Control analysis in which the AUC summary of multi-resolution partition similarity was computed considering only the sections of the parameter space for which empirical partitions outperformed the spin null model.

**FIG. S3.**
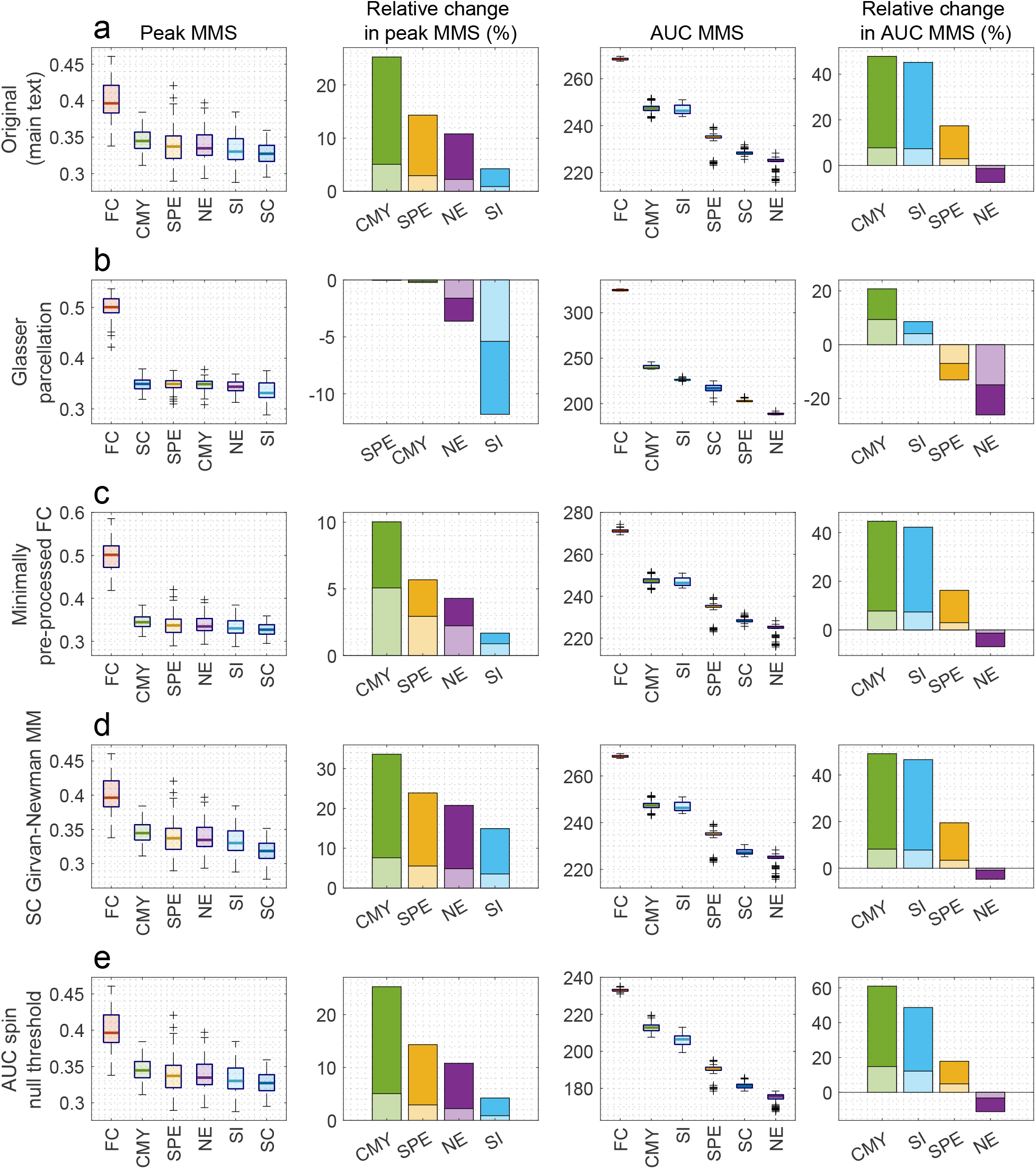
Control analyses using the MMS partition similarity measure. To facilitate the comparison of control analyses to the results presented in the main text, panel **(a)** recapitulates the results shown in Fig. 3b,c,d,e (respectively, left to right). The main text analysis was conducted considering (i) the Schaefer 200 parcellation, (ii) fMRI data pre-processed to regress out the mean global signal and other nuisance variables, (iii) SC matrices clustered using the Potts null model for modularity maximization, and (iv) the AUC summary of multi-resolution partition similarity computed across the entire resolution parameter space. In each control analysis, we tested the robustness of our results to changes in one of these four factors, while the other factors remained unaltered. **(b)** Control analysis using the Glasser parcellation. **(c)** Control analysis using the ICA FIX minimally preprocessed fMRI data from the Human Connectome Project, with no additional processing steps. **(d)** Control analysis using the Girvan–Newman null model for the modularity maximization of SC matrices. **(e)** Control analysis in which the AUC summary of multi-resolution partition similarity was computed considering only the sections of the parameter space for which empirical partitions outperformed the spin null model.

## References

[1] D. S. Bassett and O. Sporns, “Network neuroscience,” Nat. Neurosci., vol. 20, pp. 353–364, Feb. 2017.

[2] A. Fornito, A. Zalesky, and E. T. Bullmore, Fundamentals of brain network analysis. 2016.

[3] R. C. Craddock, R. Cameron Craddock, S. Jbabdi, C.-G. Yan, J. T. Vogelstein, F. Xavier Castellanos, A. Di Martino, C. Kelly, K. Heberlein, S. Colcombe, and M. P. Milham, “Imaging human connectomes at the macroscale,” 2013.

[4] H.-J. Park and K. Friston, “Structural and functional brain networks: from connections to cognition,” Science, vol. 342, p. 1238411, Nov. 2013.

[5] J. S. Damoiseaux and M. D. Greicius, “Greater than the sum of its parts: a review of studies combining structural connectivity and resting-state functional connectivity,” Brain Struct. Funct., vol. 213, pp. 525–533, Oct. 2009.

[6] L. E. Súarez, R. D. Markello, R. F. Betzel, and B. Misic, “Linking structure and function in macroscale brain networks,” Trends Cogn. Sci., vol. 24, pp. 302–315, Apr. 2020.

[7] O. Sporns and R. F. Betzel, “Modular brain networks,” Annu. Rev. Psychol., vol. 67, pp. 613–640, 2016.

[8] D. Meunier, R. Lambiotte, and E. T. Bullmore, “Modular and hierarchically modular organization of brain networks,” Front. Neurosci., vol. 4, p. 200, 2010.

[9] R. F. Betzel, “Community detection in network neuroscience,” arXiv preprint 2011.06723, 2020.

[10] D. A. Fair, A. L. Cohen, J. D. Power, N. U. F. Dosenbach, J. A. Church, F. M. Miezin, B. L. Schlaggar, and S. E. Petersen, “Functional brain networks develop from a “local to distributed” organization,” PLoS Comput. Biol., vol. 5, p. e1000381, May 2009.

[11] M. G. Puxeddu, J. Faskowitz, R. F. Betzel, M. Petti, L. Astolfi, and O. Sporns, “The modular organization of brain cortical connectivity across the human lifespan,” Neuroimage, vol. 218, p. 116974, Sept. 2020.

[12] D. Meunier, S. Achard, A. Morcom, and E. Bullmore, “Age-related changes in modular organization of human brain functional networks,” Neuroimage, vol. 44, pp. 715–723, Feb. 2009.

[13] D. S. Bassett, N. F. Wymbs, M. A. Porter, P. J. Mucha, J. M. Carlson, and S. T. Grafton, “Dynamic reconfiguration of human brain networks during learning,” Proc. Natl. Acad. Sci. U. S. A., vol. 108, pp. 7641–7646, May 2011.

[14] M. W. Cole, J. R. Reynolds, J. D. Power, G. Repovs, A. Anticevic, and T. S. Braver, “Multi-task connectivity reveals flexible hubs for adaptive task control,” Nat. Neurosci., vol. 16, pp. 1348–1355, Sept. 2013.

[15] K. Finc, K. Bonna, X. He, D. M. Lydon-Staley, S. Kühn, W. Duch, and D. S. Bassett, “Dynamic reconfiguration of functional brain networks during working memory training,” Nat. Commun., vol. 11, p. 2435, May 2020.

[16] M. P. van den Heuvel and O. Sporns, “A cross-disorder connectome landscape of brain dysconnectivity,” Nat. Rev. Neurosci., vol. 20, pp. 435–446, July 2019.

[17] B. Mišić, R. F. Betzel, M. A. de Reus, M. P. van den Heuvel, M. G. Berman, A. R. McIntosh, and O. Sporns, “Network-Level Structure-Function relationships in human neocortex,” Cereb. Cortex, vol. 26, pp. 3285–3296, July 2016.

[18] P. Hagmann, L. Cammoun, X. Gigandet, R. Meuli, C. J. Honey, V. J. Wedeen, and O. Sporns, “Mapping the structural core of human cerebral cortex,” PLoS Biol., vol. 6, p. e159, July 2008.

[19] J. Stiso and D. S. Bassett, “Spatial embedding imposes constraints on neuronal network architectures,” Trends Cogn. Sci., vol. 22, pp. 1127–1142, Dec. 2018.

[20] J. D. Power, A. L. Cohen, S. M. Nelson, G. S. Wig, K. A. Barnes, J. A. Church, A. C. Vogel, T. O. Laumann, F. M. Miezin, B. L. Schlaggar, and S. E. Petersen, “Functional network organization of the human brain,” Neuron, vol. 72, pp. 665–678, Nov. 2011.

[21] Y. He, J. Wang, L. Wang, Z. J. Chen, C. Yan, H. Yang, H. Tang, C. Zhu, Q. Gong, Y. Zang, and A. C. Evans, “Uncovering intrinsic modular organization of spontaneous brain activity in humans,” PLoS One, vol. 4, p. e5226, Apr. 2009.

[22] B. T. T. Yeo, F. M. Krienen, J. Sepulcre, M. R. Sabuncu, D. Lashkari, M. Hollinshead, J. L. Roffman, J. W. Smoller, L. Zöllei, J. R. Polimeni, B. Fischl, H. Liu, and R. L. Buckner, “The organization of the human cerebral cortex estimated by intrinsic functional connectivity,” J. Neurophysiol., vol. 106, pp. 1125–1165, Sept. 2011.

[23] J. S. Damoiseaux, S. A. R. Rombouts, F. Barkhof, P. Scheltens, C. J. Stam, S. M. Smith, and C. F. Beckmann, “Consistent resting-state networks across healthy subjects,” 2006.

[24] N. A. Crossley, A. Mechelli, P. E. Vértes, T. T. Winton-Brown, A. X. Patel, C. E. Ginestet, P. McGuire, and E. T. Bullmore, “Cognitive relevance of the community structure of the human brain functional coactivation network,” Proc. Natl. Acad. Sci. U. S. A., vol. 110, pp. 11583–11588, July 2013.

[25] C. J. Honey, J.-P. Thivierge, and O. Sporns, “Can structure predict function in the human brain?,” 2010.

[26] M. P. van den Heuvel, R. C. W. Mandl, R. S. Kahn, and H. E. Hulshoff Pol, “Functionally linked resting-state networks reflect the underlying structural connectivity architecture of the human brain,” Hum. Brain Mapp., vol. 30, pp. 3127–3141, Oct. 2009.

[27] P. N. Alves, C. Foulon, V. Karolis, D. Bzdok, D. S. Margulies, E. Volle, and M. Thiebaut de Schotten, “An improved neuroanatomical model of the default-mode network reconciles previous neuroimaging and neuropathological findings,” Commun Biol, vol. 2, p. 370, Oct. 2019.

[28] A. Avena-Koenigsberger, B. Misic, and O. Sporns, “Communication dynamics in complex brain networks,” Nat. Rev. Neurosci., vol. 19, pp. 17–33, Dec. 2018.

[29] D. Graham, A. Avena-Koenigsberger, and B. Mišić, “Editorial: Network communication in the brain,” Netw Neurosci, vol. 4, pp. 976–979, Nov. 2020.

[30] C. Seguin, Y. Tian, and A. Zalesky, “Network communication models improve the behavioral and functional predictive utility of the human structural connectome,” Network Neuroscience, 2020.

[31] J. Goñi, M. P. van den Heuvel, A. Avena-Koenigsberger, N. Velez de Mendizabal, R. F. Betzel, A. Griffa, P. Hagmann, B. Corominas-Murtra, J.-P. Thiran, and O. Sporns, “Resting-brain functional connectivity predicted by analytic measures of network communication,” Proc. Natl. Acad. Sci. U. S. A., vol. 111, pp. 833–838, Jan. 2014.

[32] C. Seguin, A. Razi, and A. Zalesky, “Inferring neural signalling directionality from undirected structural connectomes,” Nat. Commun., vol. 10, p. 4289, Sept. 2019.

[33] M. Kaiser and C. C. Hilgetag, “Nonoptimal component placement, but short processing paths, due to long-distance projections in neural systems,” PLoS Comput. Biol., vol. 2, p. e95, July 2006.

[34] V. Latora and M. Marchiori, “Efficient behavior of small-world networks,” Phys. Rev. Lett., vol. 87, p. 198701, Nov. 2001.

[35] M. Boguna, D. Krioukov, and K. C. Claffy, “Navigability of complex networks,” Nat. Phys., vol. 5, no. 1, pp. 74–80, 2009.

[36] C. Seguin, M. P. van den Heuvel, and A. Zalesky, “Navigation of brain networks,” Proc. Natl. Acad. Sci. U. S. A., vol. 115, pp. 6297–6302, June 2018.

[37] M. Rosvall, A. Grönlund, P. Minnhagen, and K. Sneppen, “Searchability of networks,” Phys. Rev. E Stat. Nonlin. Soft Matter Phys., vol. 72, p. 046117, Oct. 2005.

[38] E. Estrada and N. Hatano, “Communicability in complex networks,” Phys. Rev. E Stat. Nonlin. Soft Matter Phys., vol. 77, p. 036111, Mar. 2008.

[39] J. Andreotti, K. Jann, L. Melie-Garcia, S. Giezendanner, E. Abela, R. Wiest, T. Dierks, and A. Federspiel, “Validation of network communicability metrics for the analysis of brain structural networks,” PLoS One, vol. 9, no. 12, p. e115503, 2014.

[40] D. C. Van Essen, S. M. Smith, D. M. Barch, T. E. J. Behrens, E. Yacoub, K. Ugurbil, and WU-Minn HCP Consortium, “The WU-Minn human connectome project: an overview,” Neuroimage, vol. 80, pp. 62–79, Oct. 2013.

[41] A. Schaefer, R. Kong, E. M. Gordon, T. O. Laumann, X.-N. Zuo, A. J. Holmes, S. B. Eickhoff, and B. T. T. Yeo, “Local-Global parcellation of the human cerebral cortex from intrinsic functional connectivity MRI,” Cereb. Cortex, vol. 28, pp. 3095–3114, Sept. 2018.

[42] P. E. Vértes, A. F. Alexander-Bloch, N. Gogtay, J. N. Giedd, J. L. Rapoport, and E. T. Bullmore, “Simple models of human brain functional networks,” Proc. Natl. Acad. Sci. U. S. A., vol. 109, pp. 5868–5873, Apr. 2012.

[43] R. F. Betzel, A. Avena-Koenigsberger, J. Goñi, Y. He, M. A. de Reus, A. Griffa, P. E. Vértes, B. Mišic, J.-P. Thiran, P. Hagmann, M. van den Heuvel, X.-N. Zuo, E. T. Bullmore, and O. Sporns, “Generative models of the human connectome,” Neuroimage, vol. 124, pp. 1054–1064, Jan. 2016.

[44] K. Shen, A. Goulas, D. S. Grayson, J. Eusebio, J. S. Gati, R. S. Menon, A. R. McIntosh, and S. Everling, “Exploring the limits of network topology estimation using diffusion-based tractography and tracer studies in the macaque cortex,” Neuroimage, vol. 191, pp. 81–92, May 2019.

[45] D. S. Margulies, S. S. Ghosh, A. Goulas, M. Falkiewicz, J. M. Huntenburg, G. Langs, G. Bezgin, S. B. Eickhoff, F. X. Castellanos, M. Petrides, E. Jefferies, and J. Smallwood, “Situating the default-mode network along a principal gradient of macroscale cortical organization,” Proc. Natl. Acad. Sci. U. S. A., vol. 113, pp. 12574–12579, Nov. 2016.

[46] B. Vézquez-Rodríguez, Z.-Q. Liu, P. Hagmann, and B. Misic, “Signal propagation via cortical hierarchies,” Network Neuroscience, vol. 4, pp. 1072–1090, Nov. 2020.

[47] P. Wang, R. Kong, X. Kong, R. Liégeois, C. Orban, G. Deco, M. P. van den Heuvel, and B. T. Thomas Yeo, “Inversion of a large-scale circuit model reveals a cortical hierarchy in the dynamic resting human brain,” Sci Adv, vol. 5, p. eaat7854, Jan. 2019.

[48] C. Paquola, R. Vos De Wael, K. Wagstyl, R. A. I. Bethlehem, S.-J. Hong, J. Seidlitz, E. T. Bullmore, A. C. Evans, B. Misic, D. S. Margulies, J. Smallwood, and B. C. Bernhardt, “Microstructural and functional gradients are increasingly dissociated in transmodal cor-tices,” PLoS Biol., vol. 17, p. e3000284, May 2019.

[49] E. Dhamala, K. W. Jamison, A. Jaywant, S. Dennis, and A. Kuceyeski, “Distinct functional and structural connections predict crystallised and fluid cognition in healthy adults,” Hum. Brain Mapp., vol. 42, pp. 3102–3118, July 2021.

[50] R. H. Kaiser, J. R. Andrews-Hanna, T. D. Wager, and D. A. Pizzagalli, “Large-Scale network dysfunction in major depressive disorder,” 2015.

[51] L. Q. Uddin, B. T. T. Yeo, and R. N. Spreng, “Towards a universal taxonomy of macro-scale functional human brain networks,” Brain Topogr., vol. 32, pp. 926–942, Nov. 2019.

[52] L. G. S. Jeub, O. Sporns, and S. Fortunato, “Multiresolution consensus clustering in networks,” Sci. Rep., vol. 8, p. 3259, Feb. 2018.

[53] J. A. Contreras, A. Avena-Koenigsberger, S. L. Risacher, J. D. West, E. Tallman, B. C. McDonald, M. R. Farlow, L. G. Apostolova, J. Goñi, M. Dzemidzic, Y.-C. Wu, D. Kessler, L. Jeub, S. Fortunato, A. J. Saykin, and O. Sporns, “Resting state network modularity along the prodromal late onset alzheimer’s disease continuum,” Neuroimage Clin, vol. 22, p. 101687, Jan. 2019.

[54] Y. He, S. Lim, S. Fortunato, O. Sporns, L. Zhang, J. Qiu, P. Xie, and X.-N. Zuo, “Reconfiguration of cortical networks in MDD uncovered by multiscale community detection with fMRI,” Cereb. Cortex, vol. 28, pp. 1383–1395, Apr. 2018.

[55] V. D. Blondel, J.-L. Guillaume, R. Lambiotte, and E. Lefebvre, “Fast unfolding of communities in large networks,” J. Stat. Mech: Theory Exp., 2008.

[56] S. Fortunato and D. Hric, “Community detection in networks: A user guide,” Phys. Rep., vol. 659, pp. 1–44, 2016.

[57] N. X. Vinh, J. Epps, and J. Bailey, “Information theoretic measures for clusterings comparison,” in Proceedings of the 26th Annual International Conference on Machine Learning - ICML ‘09, (New York, New York, USA), ACM Press, 2009.

[58] A. F. Alexander-Bloch, H. Shou, S. Liu, T. D. Satterthwaite, D. C. Glahn, R. T. Shinohara, S. N. Vandekar, and A. Raznahan, “On testing for spatial correspondence between maps of human brain structure and function,” Neuroimage, vol. 178, pp. 540–551, Sept. 2018.

[59] F. Vaša, J. Seidlitz, R. Romero-Garcia, K. J. Whitaker, G. Rosenthal, P. E. Vértes, M. Shinn, A. Alexander-Bloch, P. Fonagy, R. J. Dolan, P. B. Jones, I. M. Goodyer, NSPN consortium, O. Sporns, and E. T. Bullmore, “Adolescent tuning of association cortex in human structural brain networks,” Cereb. Cortex, vol. 28, pp. 281–294, Jan. 2018.

[60] R. D. Markello and B. Misic, “Comparing spatial null models for brain maps,” Neuroimage, vol. 236, p. 118052, Aug. 2021.

[61] R. F. Betzel, A. Griffa, A. Avena-Koenigsberger, J. Goñi, J.-P. Thiran, P. Hagmann, and O. Sporns, “Multi-scale community organization of the human structural connectome and its relationship with restingstate functional connectivity,” Network Science, vol. 1, no. 3, pp. 353–373, 2013.

[62] B. Vázquez-Rodríguez, L. E. Súarez, R. D. Markello, G. Shafiei, C. Paquola, P. Hagmann, M. P. van den Heuvel, B. C. Bernhardt, R. N. Spreng, and B. Misic, “Gradients of structure-function tethering across neocortex,” Proc. Natl. Acad. Sci. U. S. A., vol. 116, pp. 21219–21227, Oct. 2019.

[63] F. Z. Esfahlani, J. Faskowitz, J. Slack, B. Mišić, and R. F. Betzel, “Local structure-function relationships in human brain networks across the lifespan.”

[64] A. Avena-Koenigsberger, X. Yan, A. Kolchinsky, M. van den Heuvel, P. Hagmann, and O. Sporns, “A spectrum of routing strategies for brain networks,” PLoS Comput. Biol., vol. 15, p. e1006833, Mar. 2019.

[65] J. Goñi, A. Avena-Koenigsberger, N. Velez de Mendizabal, M. P. van den Heuvel, R. F. Betzel, and O. Sporns, “Exploring the morphospace of communication efficiency in complex networks,” PLoS One, vol. 8, no. 3, p. e58070, 2013.

[66] E. Estrada, M. Benzi, and N. Hatano, The Physics of Communicability in Complex Networks. 2012.

[67] Y. Osmanlioğlu, B. Tunç, D. Parker, M. A. Elliott, G. L. Baum, R. Ciric, T. D. Satterthwaite, R. E. Gur, R. C. Gur, and R. Verma, “System-level matching of structural and functional connectomes in the human brain,” Neuroimage, vol. 199, pp. 93–104, Oct. 2019.

[68] M. Rosvall and C. T. Bergstrom, “Maps of random walks on complex networks reveal community structure,” Proc Natl Acad Sci U S A, vol. 105, pp. 1118–23, Jan 2008.

[69] J. J. Crofts, D. J. Higham, R. Bosnell, S. Jbabdi, P. M. Matthews, T. E. J. Behrens, and H. Johansen-Berg, “Network analysis detects changes in the contralesional hemisphere following stroke,” Neuroimage, vol. 54, pp. 161–169, Jan. 2011.

[70] E. Lella and E. Estrada, “Communicability distance reveals hidden patterns of alzheimer’s disease,” 2020.

[71] J. M. Shine, M. J. Aburn, M. Breakspear, and R. A. Poldrack, “The modulation of neural gain facilitates a transition between functional segregation and integration in the brain,” Elife, vol. 7, Jan. 2018.

[72] D. S. Grayson, E. Bliss-Moreau, C. J. Machado, J. Bennett, K. Shen, K. A. Grant, D. A. Fair, and D. G. Amaral, “The rhesus monkey connectome predicts disrupted functional networks resulting from pharmacogenetic inactivation of the amygdala,” Neuron, vol. 91, pp. 453–466, July 2016.

[73] R. F. Betzel, J. D. Medaglia, A. E. Kahn, J. Soffer, D. R. Schonhaut, and D. S. Bassett, “Structural, geometric and genetic factors predict interregional brain connectivity patterns probed by electrocorticography,” Nat Biomed Eng, vol. 3, pp. 902–916, Nov. 2019.

[74] F. Zamani Esfahlani, Y. Jo, M. G. Puxeddu, H. Merritt, J. C. Tanner, S. Greenwell, R. Patel, J. Faskowitz, and R. F. Betzel, “Modularity maximization as a flexible and generic framework for brain network exploratory analysis,” Neuroimage, vol. 244, p. 118607, Dec. 2021.

[75] R. F. Betzel, J. D. Medaglia, L. Papadopoulos, G. L. Baum, R. Gur, R. Gur, D. Roalf, T. D. Satterthwaite, and D. S. Bassett, “The modular organization of human anatomical brain networks: Accounting for the cost of wiring,” Netw Neurosci, vol. 1, pp. 42–68, Feb. 2017.

[76] M. G. Puxeddu, J. Faskowitz, O. Sporns, L. Astolfi, and R. F. Betzel, “Multi-modal and multi-subject modular organization of human brain networks.” Jan. 2022.

[77] I. Diez, P. Bonifazi, I. Escudero, B. Mateos, M. A. Muñoz, S. Stramaglia, and J. M. Cortes, “A novel brain partition highlights the modular skeleton shared by structure and function,” Sci. Rep., vol. 5, p. 10532, June 2015.

[78] R. F. Betzel, J. D. Medaglia, and D. S. Bassett, “Diversity of meso-scale architecture in human and non-human connectomes,” Nat. Commun., vol. 9, p. 346, Jan. 2018.

[79] J. Faskowitz, X. Yan, X.-N. Zuo, and O. Sporns, “Weighted stochastic block models of the human connectome across the life span,” Sci. Rep., vol. 8, p. 12997, Aug. 2018.

[80] G. Deco, A. R. McIntosh, K. Shen, R. M. Hutchison, R. S. Menon, S. Everling, P. Hagmann, and V. K. Jirsa, “Identification of optimal structural connectivity using functional connectivity and neural modeling,” J. Neurosci., vol. 34, pp. 7910–7916, June 2014.

[81] A. Messé, D. Rudrauf, A. Giron, and G. Marrelec, “Predicting functional connectivity from structural connectivity via computational models using MRI: an extensive comparison study,” Neuroimage, vol. 111, pp. 65–75, May 2015.

[82] C.-H. Yeh, R. E. Smith, X. Liang, F. Calamante, and A. Connelly, “Correction for diffusion MRI fibre tracking biases: The consequences for structural connectomic metrics,” Neuroimage, vol. 142, pp. 150–162, Nov. 2016.

[83] F. Zhang, A. Daducci, Y. He, S. Schiavi, C. Seguin, R. E. Smith, C.-H. Yeh, T. Zhao, and L. J. O’Donnell, “Quantitative mapping of the brain’s structural connectivity using diffusion MRI tractography: A review,” Neuroimage, vol. 249, p. 118870, Jan. 2022.

[84] A. Zalesky, A. Fornito, L. Cocchi, L. L. Gollo, M. P. van den Heuvel, and M. Breakspear, “Connectome sensitivity or specificity: which is more important?,” Neuroimage, June 2016.

[85] M. F. Glasser, S. N. Sotiropoulos, J. A. Wilson, T. S. Coalson, B. Fischl, J. L. Andersson, J. Xu, S. Jbabdi, M. Webster, J. R. Polimeni, D. C. Van Essen, M. Jenkinson, and WU-Minn HCP Consortium, “The minimal preprocessing pipelines for the human connectome project,” Neuroimage, vol. 80, pp. 105–124, Oct. 2013.

[86] S. N. Sotiropoulos, S. Jbabdi, J. Xu, J. L. Andersson, S. Moeller, E. J. Auerbach, M. F. Glasser, M. Hernandez, G. Sapiro, M. Jenkinson, D. A. Feinberg, E. Yacoub, C. Lenglet, D. C. Van Essen, K. Ugurbil, T. E. J. Behrens, and WU-Minn HCP Consortium, “Advances in diffusion MRI acquisition and processing in the human connectome project,” Neuroimage, vol. 80, pp. 125–143, Oct. 2013.

[87] J.-D. Tournier, F. Calamante, and A. Connelly, “MR-trix: Diffusion tractography in crossing fiber regions,” Int. J. Imaging Syst. Technol., vol. 22, Mar. 2012.

[88] J.-D. Tournier, F. Calamante, and A. Connelly, “Robust determination of the fibre orientation distribution in diffusion MRI: non-negativity constrained superresolved spherical deconvolution,” Neuroimage, vol. 35, pp. 1459–1472, May 2007.

[89] J. D. Tournier, F. Calamante, and A. Connelly, “Improved probabilistic streamlines tractography by 2nd order integration over fibre orientation distributions,” in Proceedings of the international society for magnetic resonance in medicine, vol. 18, p. 1670, 2010.

[90] R. E. Smith, J.-D. Tournier, F. Calamante, and A. Connelly, “Anatomically-constrained tractography: improved diffusion MRI streamlines tractography through effective use of anatomical information,” Neuroimage, vol. 62, pp. 1924–1938, Sept. 2012.

[91] S. Mansour L, Y. Tian, B. T. T. Yeo, V. Cropley, and A. Zalesky, “High-resolution connectomic fingerprints: Mapping neural identity and behavior,” Neuroimage, vol. 229, p. 117695, Apr. 2021.

[92] M. F. Glasser, T. S. Coalson, E. C. Robinson, C. D. Hacker, J. Harwell, E. Yacoub, K. Ugurbil, J. Andersson, C. F. Beckmann, M. Jenkinson, S. M. Smith, and D. C. Van Essen, “A multi-modal parcellation of human cerebral cortex,” Nature, vol. 536, pp. 171–178, Aug. 2016.

[93] R. F. Betzel, A. Griffa, P. Hagmann, and B. Mišić, “Distance-dependent consensus thresholds for generating group-representative structural brain networks,” Netw Neurosci, vol. 3, no. 2, pp. 475–496, 2019.

[94] S. M. Smith, C. F. Beckmann, J. Andersson, E. J. Auerbach, J. Bijsterbosch, G. Douaud, E. Duff, D. A. Feinberg, L. Griffanti, M. P. Harms, M. Kelly, T. Laumann, K. L. Miller, S. Moeller, S. Petersen, J. Power, G. Salimi-Khorshidi, A. Z. Snyder, A. T. Vu, M. W. Woolrich, J. Xu, E. Yacoub, K. Uğurbil, D. C. Van Essen, M. F. Glasser, and WU-Minn HCP Consortium, “Resting-state fMRI in the human connectome project,” Neuroimage, vol. 80, pp. 144–168, Oct. 2013.

[95] L. Parkes, B. Fulcher, M. Yücel, and A. Fornito, “An evaluation of the efficacy, reliability, and sensitivity of motion correction strategies for resting-state functional MRI,” Neuroimage, vol. 171, pp. 415–436, May 2018.

[96] T. D. Satterthwaite, M. A. Elliott, R. T. Gerraty, K. Ruparel, J. Loughead, M. E. Calkins, S. B. Eickhoff, H. Hakonarson, R. C. Gur, R. E. Gur, and D. H. Wolf, “An improved framework for confound regression and filtering for control of motion artifact in the preprocessing of resting-state functional connectivity data,” Neuroimage, vol. 64, pp. 240–256, Jan. 2013.

[97] M. Rubinov and O. Sporns, “Complex network measures of brain connectivity: uses and interpretations,” Neuroimage, vol. 52, pp. 1059–1069, Sept. 2010.

[98] A. Avena-Koenigsberger, B. Mišić, R. X. D. Hawkins, A. Griffa, P. Hagmann, J. Goñi, and O. Sporns, “Path ensembles and a tradeoff between communication efficiency and resilience in the human connectome,” Brain Struct. Funct., pp. 1–16, 2016.

[99] J. J. Crofts and D. J. Higham, “A weighted communicability measure applied to complex brain networks,” J. R. Soc. Interface, vol. 6, pp. 411–414, Apr. 2009.

[100] V. A. Traag, P. Van Dooren, and Y. Nesterov, “Narrow scope for resolution-limit-free community detection,” Phys. Rev. E Stat. Nonlin. Soft Matter Phys., vol. 84, p. 016114, July 2011.

[101] M. Bazzi, M. A. Porter, S. Williams, M. McDonald, D. J. Fenn, and S. D. Howison, “Community detection in temporal multilayer networks, with an application to correlation networks,” 2016.

[102] M. Girvan and M. E. J. Newman, “Community structure in social and biological networks,” Proc. Natl. Acad. Sci. U. S. A., vol. 99, pp. 7821–7826, June 2002.

[103] M. E. J. Newman and M. Girvan, “Finding and evaluating community structure in networks,” Phys. Rev. E Stat. Nonlin. Soft Matter Phys., vol. 69, p. 026113, Feb. 2004.

[104] “Website.” XuanVinhNguyen (2022). TheAdjustedMutualInformation (https://www.mathworks.com/matlabcentral/fileexchange/33144-the-adjusted-mutual-information),MATLABCentralFileExchange.RetrievedJanuary26,2022.

[105] J. Munkres, “Algorithms for the assignment and transportation problems,” 1957.

[106] “Website.” YiCao (2022). HungarianAlgorithmforLinearAssignmentProblems (V2. 3)(https://www.mathworks.com/matlabcentral/fileexchange/20652-hungarian-algorithm-for-linear-assignment-problems-,MATLABCentralFileExchange.RetrievedJanuary26,2022.

